# Inducible and reversible inhibition of miRNA-mediated gene repression *in vivo*

**DOI:** 10.1101/2021.06.01.445680

**Authors:** Gaspare La Rocca, Bryan King, Bing Shui, Xiaoyi Li, Minsi Zhang, Kemal Akat, Paul Ogrodowski, Chiara Mastroleo, Kevin Chen, Vincenzo Cavalieri, Yilun Ma, Viviana Anelli, Doron Betel, Joana A. Vidigal, Thomas Tuschl, Gunter Meister, Craig B. Thompson, Tullia Lindsten, Kevin M. Haigis, Andrea Ventura

## Abstract

Although virtually all gene networks are predicted to be controlled by miRNAs, the contribution of this important layer of gene regulation to tissue homeostasis in adult animals remains unclear. Gain and loss of function experiments have provided key insights into the specific function of individual miRNAs, but effective genetic tools to study the functional consequences of global inhibition of miRNA activity *in vivo* are lacking. Here we report the generation and characterization of a genetically engineered mouse strain in which miRNA-mediated gene repression can be reversibly inhibited without affecting miRNA biogenesis or abundance. We demonstrate the usefulness of this strategy by investigating the consequences of acute inhibition of miRNA function in adult animals. We find that different tissues and organs respond differently to global loss of miRNA function. While miRNA-mediated gene repression is essential for the homeostasis of the heart and the skeletal muscle, it is largely dispensable in the majority of other organs. Even in tissues where it is not required for homeostasis, such as the intestine and hematopoietic system, miRNA activity can become essential during regeneration following acute injury. These data support a model where many metazoan tissues primarily rely on miRNA function to respond to potentially pathogenic events.

---

MicroRNAs (miRNAs) are short non-coding RNAs that in Metazoa repress gene expression at the post-transcriptional level by binding to partially complementary sequences on target mRNAs^1–4^.

MiRNAs act as part of a large ribonucleoprotein complex known as the miRNA-Induced Silencing Complex (miRISC). In mammals, the Argonaute protein family (AGO1-4) and the Trinucleotide Repeat-Containing gene 6 protein family (TNRC6A/GW182, TNRC6B and TNRC6C) are the core components of the miRISC. AGO binds to the miRNA, and facilitates its interaction with target mRNAs^5^. In turn, TNRC6 binds to AGO and recruits the decapping and deadenlyation complexes, leading to degradation of target mRNAs^6–16^.

Although miRNAs are abundantly expressed in embryonic and adult mouse tissues, and computational and experimental analyses indicate that they target components of virtually every cellular process^17^, animals harboring targeted deletion of single miRNA genes are often indistinguishable from their wild type counterparts^18–25^. One explanation for these observations is that the redundant functions of related miRNAs may buffer the emergence of obvious phenotypes in mutant animals^1, 3^. Interestingly, however, clear phenotypes often emerge in mutant adult animals when exposed to external or internal perturbations^19, 23, 26^. These observations suggest that, at least in some contexts, miRNA function is conditionally, rather than constitutively, required to carry on cellular processes.

Previous efforts to investigate the consequences of global inhibition of miRNA function have relied upon the targeted deletion of the core miRNA biogenesis factors DICER, DROSHA, and DGCR8 (reviewed in ref. ^27^). Several animal models harboring conditional or constitutive knockout alleles of these genes have been generated^28–36^. Although these strategies have provided important insights into miRNA biology, they suffer from several limitations.

First, inactivation of these gene products is known to have other consequences in addition to impairing miRNA biogenesis. For instance, DICER is involved in epigenetic regulation in the nucleus in a miRNA-independent manner^37–42^, and is essential to metabolize transcripts from short interspersed nuclear elements, predominantly Alu RNAs in humans and B1 and B2 RNAs in rodents^43^. DROSHA, on the other hand, regulates the expression of several coding and non-coding RNAs by directly cleaving stem–loop structures embedded within the transcripts^44^. Furthermore, DICER and DROSHA are also involved in ribosomal RNA biogenesis^45^ and in the DNA-damage response^46 47^, and DGCR8 regulates the maturation of small nucleolar RNAs and of some long non-coding RNAs^48, 49^. Consequently, the phenotypes observed in these models cannot be solely attributed to inhibition of miRNA activity.

Another limitation of conditional ablation of miRNA-biogenesis genes *in vivo* is that due to their high stability mature miRNAs can persist for several days after their biogenesis is inhibited. For example, four weeks after near complete conditional ablation of *Dicer1* in the muscle, the levels of the highest expressed miRNAs were found to be only reduced by 30-40% and their expression remained substantial even 18 months later^24^. This complicates the interpretation of experiments based on temporally controlled conditional ablation of these biogenesis factors, especially in non-proliferating tissues.

Third, a subset of mammalian miRNAs does not rely on the canonical biosynthesis pathway, and therefore their expression and activity are not affected by inactivation of the core miRNA biogenesis factors^44, 50–55^.

Finally, these genetic approaches are not reversible and therefore these animal models cannot be used to study the effects of transient inhibition of miRNA function.

To circumvent these limitations, we have generated a novel genetically engineered mouse strain that allows inducible and reversible disassembly of the miRISC, thereby achieving controllable inhibition of miRNA-mediated gene repression *in vivo* without affecting small RNA biogenesis. To address the reliance of adult tissues on miRNA-mediated gene repression, we have used this novel strain to investigate the consequences of acute inhibition of the miRISC under homeostatic conditions, and during tissue regeneration.

## Results

### Inhibition of the miRNA pathway through peptide-mediated disruption of the miRISC

Multiple motifs within the N-terminal domain of TNRC6 proteins contain regularly spaced tryptophan residues which mediate the interaction between AGO and TNRC6 by inserting into conserved hydrophobic pockets located on AGO’s Piwi domain^56, 57^.

A peptide encompassing one of the AGO-interacting motifs of human TNRC6B has been previously employed as an alternative to antibody-based approaches to efficiently pull down all AGO family members from cell and tissue extracts^58, 59^. This peptide, named T6B, competes with endogenous TNRC6 proteins for binding to AGOs. However, as it lacks the domains necessary for the recruitment of de-capping and de-adenylation factors, it prevents the assembly of the full miRISC, thus resulting in effective inhibition of miRISC-mediated gene repression in cells^58, 60^.

Based on these results, we reasoned that temporally and spatially controlled expression of a T6B transgene in animals would offer the unprecedented opportunity to study the consequences of acute and reversible inhibition of miRNA function *in vivo* without interfering with miRNA biogenesis or abundance (**Fig. 1a**).

**Figure 1.**
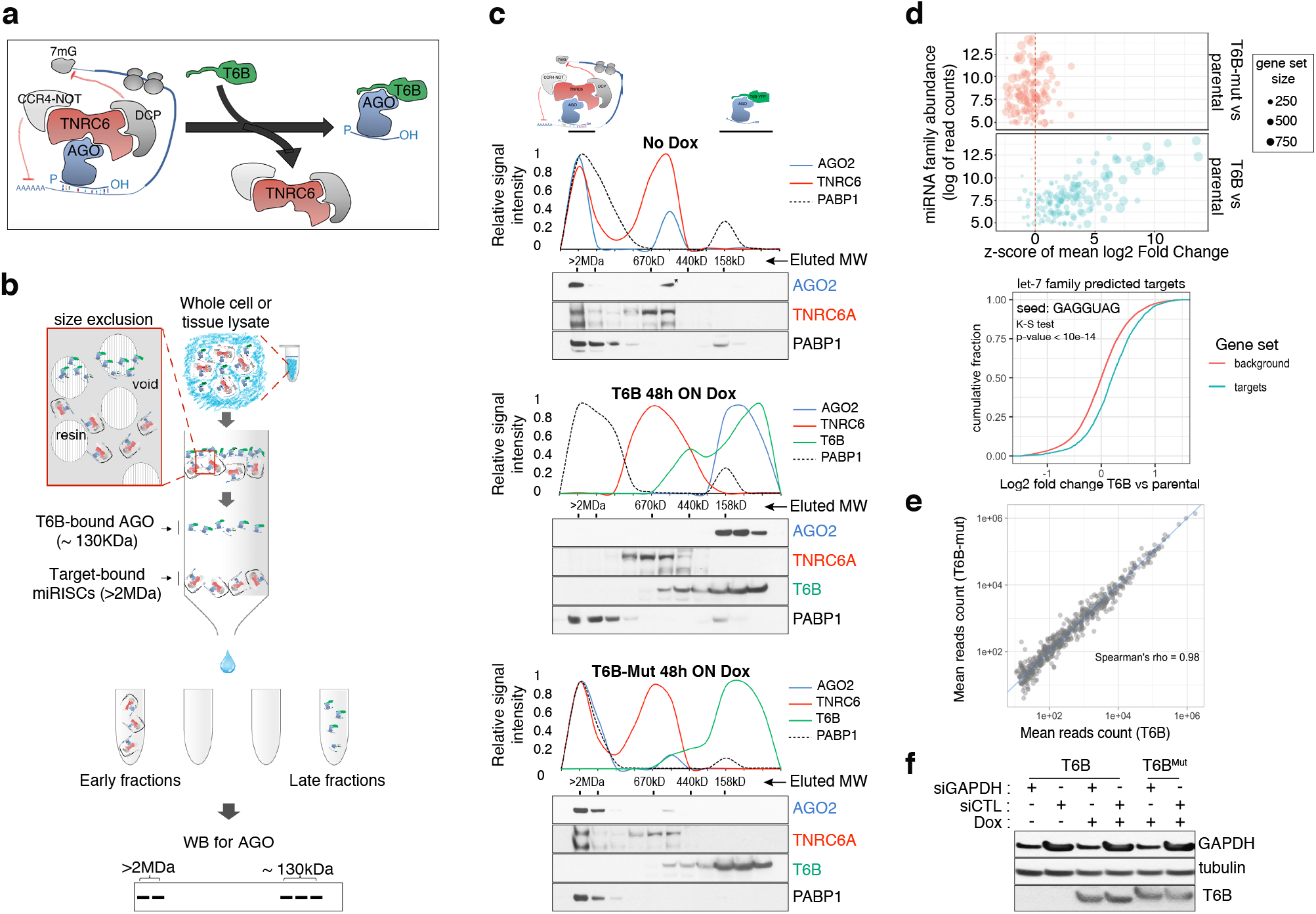
T6B fusion protein prevents miRISC assembly and impairs miRNA activity *in vitro*. **(a)** Schematics of T6B action: T6B competes with TNRC6 for binding to AGO proteins preventing miRISC assembly. **(b)** Schematics of the size exclusion chromatography (SEC) assay for the fractionation of AGO-containing complexes according to their molecular weight. **(c)** SEC profiling of miRISC components upon T6B expression: Total lysates from HCT116 cells expressing no fusion protein (upper panel), T6B (middle panel) or T6B^Mut^ (lower panel) were fractionated as described in (b) and immunoblotted to detect AGO2, TNRC6A, T6B and PABP1. **(d)** RNAseq analysis of total and small RNAs isolated from MEFs cell lines expressing either empty no fusion protein, T6B or T6B^Mut^. Upper panel: bubble plot of target derepression against miRNA abundance. The mean log2 Fold Change (T6B or T6B^Mut^ vs control) of predicted targets for each conserved miRNA family was calculated, converted to a z-score and is plotted on the x-axis against the miRNA family abundance (log of the sum of read counts for each member of the family). The size of each circle is proportional to the number of predicted targets. A positive z score indicates that the targets for that family are preferentially upregulated upon T6B expression, while a negative score would indicate preferential downregulation. Expression of T6B, but not of T6B^Mut^, causes preferential upregulation of miRNA targets of the most miRNA families and the effect is roughly proportional to each miRNA family abundance. Lower panel: cumulative distribution plot of predicted let-7 targets compared to background in T6B-expressing MEFs. **(e)** Scatter plots of miRNA abundance as determined by small-RNAseq of total RNA extracted from MEFs expressing either T6B or T6B^Mut^. Each dot represents a miRNA in miRBase. **(f)** Effect of T6B expression on AGO2 slicing activity. MEFs expressing either T6B or T6B^Mut^ were transfected with siRNAs targeting GAPDH mRNA (siGAPDH) or with scramble siRNA (siCTL). Levels of GAPDH, T6B and tubulin were assessed by immunoblot 72 hours post-transfection. T6B and T6B^Mut^ have slightly different migration on PAGE, as previously observed by Hauptmann at al.^58^.

To test the suitability of this approach, we first investigated the dynamics of interaction between T6B and the miRISC in mouse and human cell lines. We employed a previously reported size exclusion chromatography (SEC)-based assay^61, 62^ to analyze the molecular weight of AGO-containing complexes in lysates from cells expressing either a doxycycline-inducible FLAG-HA-T6B-YFP fusion protein (hereafter referred to as T6B), or a mutant version (hereafter referred to as T6B^Mut^) incapable of binding to AGO (**Extended Data Fig. 1**). We reasoned that if T6B expression prevents AGO from stably binding to TNRC6 and its targets, AGO proteins should be detected in fractions corresponding to approximately 120-130 kDa, the sum of the molecular weights of AGO (approximately 95 kDa) and the T6B fusion protein (approximately 30 kDa). In contrast, unperturbed AGO complexes that are part of the fully assembled miRISC bound to mRNAs should elute in the void of the column, which contains complexes larger than 2 MDa (**Fig. 1b**).

As expected, in lysates from cells expressing no T6B or T6B^Mut^ AGO2 and TNRC6A were mostly detected in the high molecular weight fractions, indicating the presence of target-bound miRISC (**Fig. 1c**). In contrast, AGO2 and TNRC6A were nearly completely depleted from the high molecular weight fractions in lysates from cells expressing T6B (**Fig. 1c**). Moreover, while AGO2, TNRC6A and the polyA-binding protein 1 (PABP1) cofractionated in lysates from control cells, they eluted in different fractions in lysates from T6B-expressing cells (**Fig. 1c**), indicating that T6B leads to loss of interactions between the miRISC components and mRNAs. As expected based on the strong evolutionary conservation of human and mouse AGO and TNRC6 proteins^59, 63, 64^, we obtained identical results when human T6B was expressed in mouse embryo fibroblasts (MEFs) (**Extended Data Fig. 2**).

To test whether the redistribution of AGO-containing complexes induced by T6B expression was mirrored by a loss of miRNA-mediated gene repression, we performed RNAseq analysis on MEFs expressing T6B or T6B^Mut^. Cells expressing T6B displayed marked and selective de-repression of predicted mRNA targets for expressed miRNAs (**Fig. 1d**). The extent of de-repression was roughly proportional to the abundance of individual miRNA families, with predicted targets of poorly expressed miRNAs collectively showing modest de-repression compared to targets of more abundantly expressed miRNA families (**Fig. 1d**). Importantly, de-repression of miRNA targets was not accompanied by a global change in mature miRNAs levels (**Fig. 1e)**, consistent with the role of T6B in perturbing the effector step of the miRNA pathway, without affecting miRNA processing.

Of the four mammalian AGO proteins, AGO2 is the only one that has endo-ribonucleolytic activity, which does not require TNRC6^65^ and is triggered when the AGO2-loaded small RNA and the target are perfectly complementary^66–68^. AGO2’s catalytic activity is essential for gene regulation in the germline. For example, in mouse oocytes, AGO2 loaded with endogenous small-interfering RNAs (endo-siRNAs) mediates the cleavage of coding and non-coding transcripts bearing perfectly complementary sequences^42, 69^. In metazoan somatic tissues, in contrast, AGO2 catalytic activity is mainly involved in the biogenesis of miR-486 and miR-451 in the hematopoietic system^50, 70^, and in occasional instances of miRNA-directed cleavage of mRNAs^71^.

Importantly, T6B expression does not interfere with the ability of synthetic siRNAs to cleave perfectly complementary endogenous targets (**Fig. 1f**), indicating that AGO2’s catalytic function is not affected by the binding of T6B, and implying that the loading of small RNAs onto AGOs is also not perturbed by T6B.

Collectively these results demonstrate that ectopic T6B expression in mammalian cells causes global inhibition of miRISC function with minimal perturbation of the expression of mature miRNAs, and with preservation of AGO2’s endo-nucleolytic activity.

### Generation of a mouse strain with inducible expression of a T6B transgene

To apply this general strategy to an *in vivo* setting, we next generated mouse embryonic stem cells (mESCs) expressing a doxycycline-inducible T6B transgene. We used a knock-in approach in which the doxycycline-inducible transgene is inserted into the Col1A locus of mESC expressing the reverse tetracycline-controlled transactivator (rtTA) under the control of the endogenous Rosa26 (R26) promoter^72^ (**Fig. 2a**). Targeted mESCs were tested for the ability to express the T6B transgene in response to doxycycline (**Extended Data Fig. 3**) and then used to generate mice with genotype R26^rtTA/rtTA^; ColA1^T6B/T6B^ (hereafter R26^T6B^). R26^rtTA/rtTA^; ColA1^+/+^ mice, with untargeted ColA1 loci but expressing rtTA served as negative controls (hereafter R26^CTL^).

**Figure 2.**
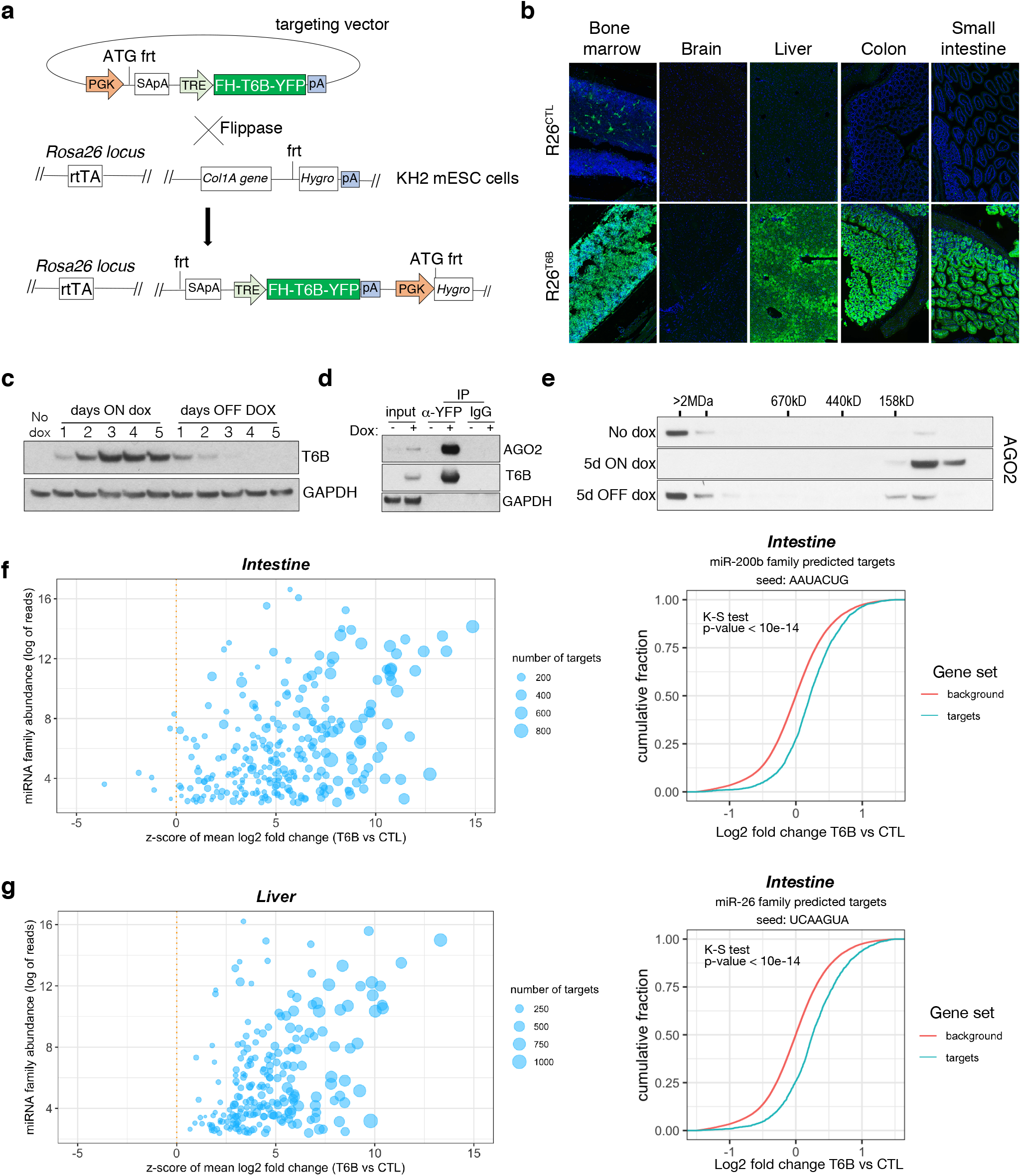
Expression of T6B reversibly blocks miRISC assembly and inhibits miRNA function *in vivo*. **(a)** Schematic of the targeting strategy to generate the T6B mouse. The construct contains a flippase recognition target site (frt) that allows homing into the Col1A locus when electroporated together with a vector expressing the Flippase recombinase into KH2 (Col1A-frt/Rosa26-rtTA) murine embryonic stem cells. KH2 also express the rtTA trans-activator driven by the endogenous Rosa26 (R26) promoter. **(b)** Immunofluorescence imaging performed using an anti-YFP antibody, showing T6B expression in a panel of tissues of adult R26^T6B^ mice fed doxycycline for 7 days. Tissues from R26^CTL^ (carrying the rtTA allele but not the T6B allele) were used as negative controls. **(c)** Protein lysates from the liver of R26^T6B^ mice on or off doxycycline-containing chow for the indicated number of days were resolved by SDS-PAGE and Western blotting was performed with anti-HA antibody to detect expression of the T6B transgene. **(d)** Co-IP experiments using an anti-YFP antibody showing interaction between AGO and T6B in total liver extracts from T6B mice on doxycycline containing chow. **(e)** SEC elution profile of AGO2-containing complexes in liver lysates from T6B mice euthanized at the indicated time points after doxycycline administration. Notice the shift of AGO2 from the high molecular weight fractions to the low molecular weight fractions after 5 days of doxycycline treatment and the reconstitution of the full miRISC after removal of doxycycline from the diet. **(f-g)** Total RNA extracted from the large intestine (f) and the liver (g) of R26^CTL^ and R26^T6B^ mice was subjected to RNAseq. Left panel: scatter plot showing the effect of T6B expression on targets of all miRNA families were generated as described in figure 1d. The abundance of each miRNA family was calculated using data set from Isakova et al.^100^. Right panel: Representative cumulative distribution plot of log2 fold changes in expression of predicted targets of the indicated miRNA families.

Upon doxycycline administration we observed strong expression of T6B in R26^T6B^ mice and across most adult tissues (**Fig. 2b**). Notable exceptions were the central nervous system (**Fig. 2b** and **Extended Data. Fig. 4**), probably due to low blood-brain barrier penetration of doxycycline, and the skeletal muscle and the heart, most likely due to low expression of the rtTA transgene in these tissues^73^.

When doxycycline was administered in the diet, T6B became detectable after 24h, reached a plateau after three days, and completely disappeared four days after doxycycline removal from the diet (**Fig. 2c**).

Because colon and liver expressed uniformly high levels of T6B in response to doxycycline, we used these tissues to test the effects of T6B expression on miRISC activity *in vivo*. Co-IP experiments using antibodies directed to T6B confirmed the interaction between AGO and T6B in these tissues (**Fig. 2d** and **Extended Data Fig. 5**). Expression of T6B resulted in nearly complete disassembly of the miRISC, as indicated by the elution shift of AGO from the high molecular weight to low molecular weight fractions in both tissues (**Fig. 2e** and **Extended Data Fig. 6**). Importantly, doxycycline removal from the diet led to a complete reconstitution of the miRISC, as indicated by the reappearance of AGO2 in the high molecular weight fractions (**Fig. 2e**).

To test whether T6B expression also resulted in inhibition of miRNA-mediated gene repression *in vivo*, we performed RNAseq on total RNAs extracted from the liver and colon of R26^T6B^ and R26^CTL^ mice kept on doxycycline-containing diet for one week. As shown in **Fig. 2f**, T6B expression resulted in marked de-repression of miRNA targets in both tissues.

Based on these results we conclude that T6B expression allows acute and reversible disruption of the miRISC, and concomitant inhibition of miRNA function *in vivo*.

### Consequences of miRISC disruption in adult tissues under homeostatic conditions

Given the central role of miRNAs in gene regulatory networks, one might expect widespread phenotypes emerging when miRISC function is systemically inhibited. Consistent with this hypothesis continuous doxycycline administration starting at conception caused embryonic lethality (**Fig. 3a**), while inhibition of miRISC starting at mid-gestation caused developmental defects and perinatal lethality in R26^T6B^ mice (**Fig. 3b** and **Extended Data Fig. 7**). Surprisingly, however, adult R26^T6B^ mice kept on doxycycline diet for up to two months remained healthy and appeared normal upon macroscopic and histopathologic examination.

**Figure 3.**
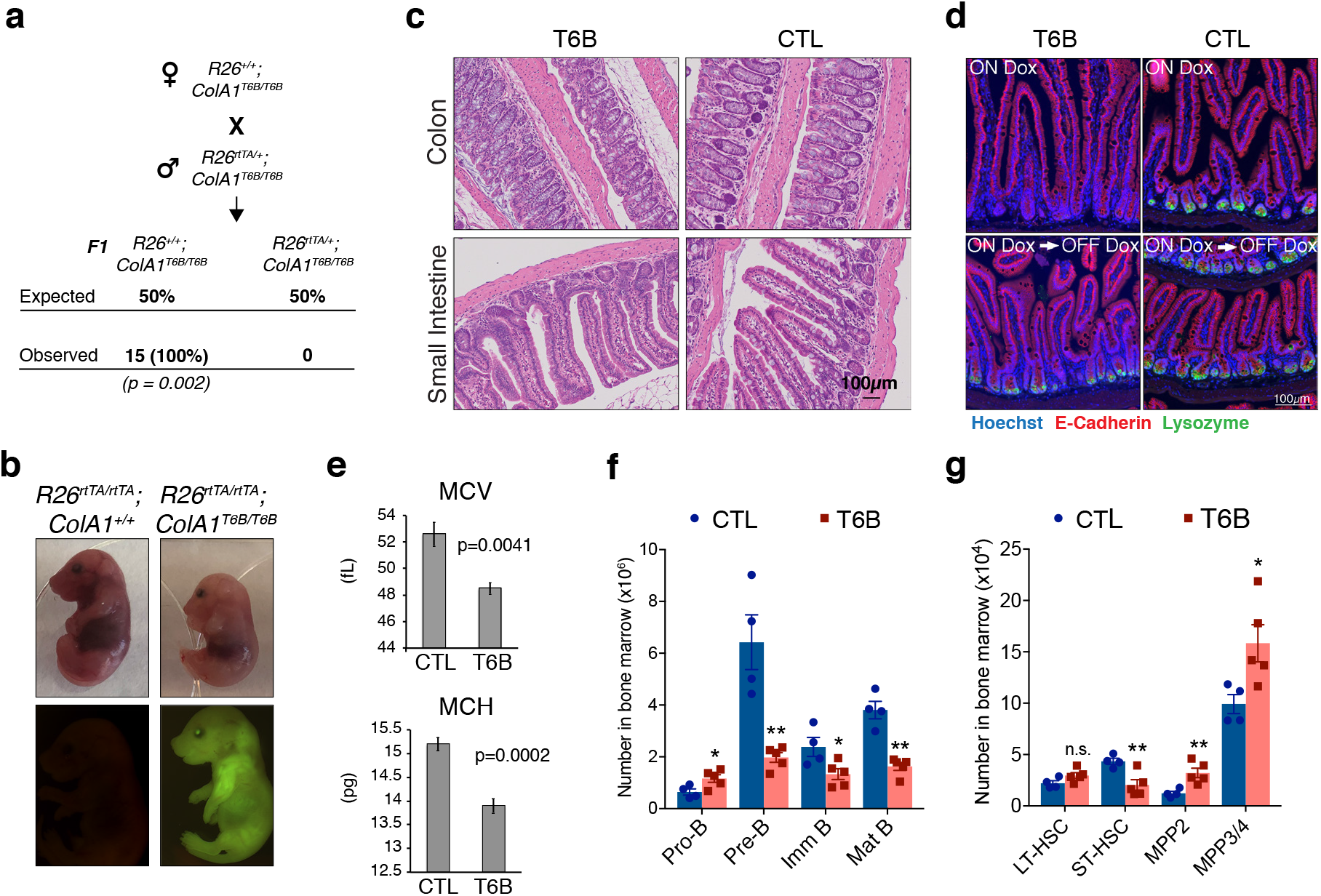
Phenotypic analysis of R26^T6B^ mice during homeostasis. **(a)** R26^+/+^; ColA1^T6B/T6B^ females were crossed with R26rt^TA/+^; ColA1^T6B/T6B^ males and doxycycline was administered by chow starting at 0.5 d.p.c. No viable pups positive for both the rtTA and T6B allele were observed (n = 15, p-value = 0.002, Fisher exact test). **(b)** Pregnant females were kept on doxycycline diet from E13.5 to E18.5 and the pups delivered on E18.5 by c-section. Note the significantly smaller size of R26^rtTA/rtTA^; ColA1^T6B/T6B^ embryos relative to R26rt^TA/rtTA^; ColA1^+/+^ control littermates. Lower row: YFP detection by epifluorescence in E18.5 pups of the indicated genotypes. **(c)** Comparison of intestine architecture in H&E sections from R26^T6B^ and R26^CTL^ mice (n = 3 for each genotype) maintained on doxycycline for 2 months. **(d)** Immunofluorescence imaging of the small intestine of R26^T6B^ and R26^CTL^ mice (n = 3-5 for each genotype) kept on doxycycline diet for a month (upper row), showing a reduction in lysozyme expression in Paneth cells in the crypts. Lysozyme expression in R26^T6B^ mice returned to normal levels upon removal of doxycycline from the diet (lower row). **(e)** Peripheral blood analysis conducted in R26^T6B^ and R26^CTL^ mice (R26^CTL^ n = 4; R26^T6B^ n = 5). **(f)** Flow cytometric analysis of bone marrow of R26^T6B^ and R26^CTL^ mice kept on doxycycline diet for 3 weeks showing developmental block at the Pro-B to Pre-B. p values (from left to right): *p = 0.0348, **p = 0.0023, *p = 0.0340, **p = 0.0004, unpaired t-test. R26^CTL^ n = 4; R26^T6B^ n = 5. **(g)** Flow cytometry analysis of the bone marrow of control and R26^T6B^ mice kept on doxycycline diet for 3 weeks. p values (from left to right): p = 0.0994, **p = 0.0092, **p = 0.0085, *p = 0.0312, unpaired t-test. R26^CTL^ n = 4; R26^T6B^ n = 5.

Detailed examination of the intestine confirmed extensive T6B expression in the epithelium and in the mesenchymal compartment (**Extended Data Fig. 8**) but no architectural abnormalities were observed (**Fig. 3c**). Cells in the crypts showed no significant changes in expression pattern of Ki67 protein (**Extended Data Fig. 9**), suggesting that the proliferation and turnover of the epithelium is maintained even in absence of a functional miRISC. No significant change in the number of goblet cells was detected throughout the intestine (**Extended Data Fig. 10**), and mice maintained normal body mass throughout the period of doxycycline treatment (**Extended Data Fig. 11**), suggesting that general intestinal functions were not affected.

Although no obvious macroscopic, functional, or architectural abnormalities were caused by T6B expression in the intestine, we observed a reduction in lysozyme expression in Paneth cells in the crypts (**Fig. 3d**, upper row). However, this phenotype was reversible, as lysozyme signal in the crypts returned to normal levels when doxycycline was removed from the diet (**Fig. 3d**, lower row), suggesting that T6B expression did not affect neither the viability of intestinal stem cells, nor their self-renewal ability.

Complete blood counts showed a modest, but significant, decrease in erythrocytes volume (MCV) and hemoglobin content (MCH) in R26^T6B^ RBCs (**Fig. 3e** and **table 1**), analogously to what reported in mice harboring targeted deletion of miR-451^74^. Flow cytometric analysis of bone marrow showed a 3-fold depletion in Pre-B cells as well as a significant decrease in immature and mature circulating B cells in R26^T6B^ mice. We also observed a reciprocal increase in the frequency of Pro-B cells in the bone marrow of these animals (**Fig. 3f** and **Extended Data Fig. 12**). These results are reminiscent of the partial block in B cell differentiation observed upon deletion of the miR-17~92 cluster^75^.

Further characterization of hematopoietic stem cells showed that the number of long-term repopulating hematopoietic stem cells (LT-HSC) was unaffected after 3 weeks of doxycycline exposure. However, we observed a modest decrease in short-term repopulating HSCs (ST-HSCs) and a concomitant increase in multipotent progenitors (MPPs) relative to controls (**Fig. 3g** and **Extended Data Fig. 13**).

Collectively, these data suggest that in a subset of adult tissues miRISC function can be suppressed with minimal or no consequences on the ability of these tissues to maintain homeostasis.

### miRISC disruption impairs the regeneration of injured colon epithelium

Several studies have shown that the phenotype caused by targeted deletion of individual miRNAs often manifests only after the mutant animals are subjected to “stress”^19, 23, 26, 76^. For example, ablation of miR-143/145 causes no apparent phenotype under homeostasis but severely impairs the ability of the mutant animals to respond to acute damage to the intestinal epithelium^19^.

Prompted by these reports, and by our initial observation that prolonged T6B expression does not substantially affect intestinal homeostasis, we tested the consequences of miRISC disruption on the regenerating intestine. A cohort of R26^T6B^ and R26^CTL^ mice were kept on doxycycline-containing diet for ten days, after which they were treated with dextran sulfate sodium (DSS), which induces severe colitis in mice^19, 77^.

A significant and progressive loss of body mass was observed in both groups during DSS treatment and two days following DSS removal (**Fig. 4a**). However, R26^T6B^ mice lost body mass more rapidly than controls and reached critical health conditions seven days after DSS removal. Three days after DSS removal, control animals started to regain weight, reaching the initial body mass within five days after DSS removal (**Fig. 4a**). In contrast, R26^T6B^ mice failed to fully recover (**Fig. 4a**), and all reached a humane endpoint within five days after DSS removal from the diet (**Fig. 4b**). Histological analysis confirmed that DSS treatment induced the disruption of the architecture of the epithelium, and the appearance of ulcerative areas to a similar extent in both R26^T6B^ and R26^CTL^ control mice, (**Fig. 4c** and **Extended Data Fig. 14**). In contrast, although five days after DSS removal the integrity of the colonic epithelium of control mice was largely reestablished with the exception of isolated dysplastic areas (**Extended Data Fig. 15**), extensive ulcerated regions persisted in the colon of R26^T6B^ mice (**Fig. 4c**). Importantly, we observed the presence of dysplastic epithelium in R26^T6B^ mice during and after DSS treatment, indicating that miRISC disruption does not completely abolish the potential of cells to proliferate, as also confirmed by Ki67 staining (**Fig. 4d**). Therefore, we speculate that other factors, such as impaired stem cell maintenance or differentiation, may be responsible for the increased susceptibility of T6B-expressing colon to DSS treatment.

**Figure 4.**
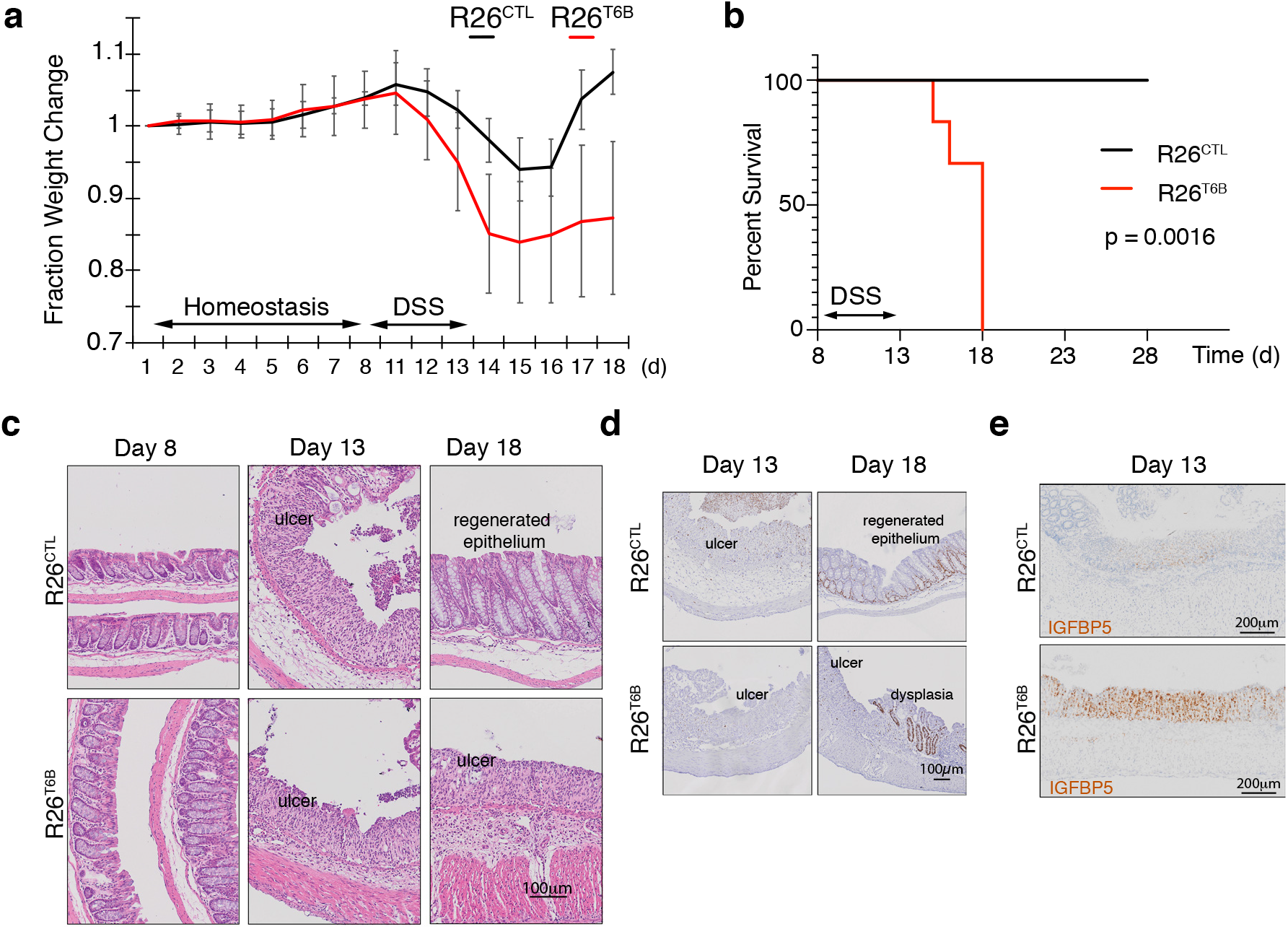
T6B-induced block of miRISC assembly leads to impaired intestinal regeneration. **(a)** R26^T6B^ and R26^CTL^ mice (n = 6 for each genotype) kept on doxycycline diet were treated with Dextran Sodium Sulphate (DSS) for 5 days to induce inflammatory colitis and their weight was monitored daily. **(b)** Kaplan-Meier curves of animals described in panel (a). **(c)** Representative hematoxylin-eosin-stained sections of intestine of R26^T6B^ and R26^CTL^ mice (n = 3 for each genotype) at different time points pre- and post-DSS treatment. **(d)** Ki67 immunostaining of section of intestine at the indicated time points. **(e)** Sections from the large intestine of control and T6B mice euthanized at day 13 were subjected to RNA in situ hybridization with a probe against the Igfbp5 transcript. The results show increased levels of IGFBP5 mRNA in ulcerated areas of R26^T6B^ as compared to controls (n = 4 for each genotype).

Chivukula and colleagues have shown that defective intestinal regeneration in the colon of miR-143/145-deficient mice is associated with upregulation of the miRNA-143 target IGFBP5 in the mesenchymal compartment. The increased levels of IGFBP5 protein cause the inhibition of IGF1R signaling in the epithelium through a non-cell autonomous mechanism, which ultimately prevented epithelial regeneration^19^. Consistent with their findings, *in situ* hybridization analyses in the colon of DSS-treated R26^T6B^ mice showed a significant upregulation of IGFBP5 mRNA in the mesenchymal compartment compared to controls (**Fig. 4e**). The extent of de-repression of IGFBP5 was comparable to that previously observed in miRNA-143/145 knockout mice^19^, providing further evidence that T6B-mediated miRISC disassembly is an effective strategy to globally inhibit miRNA function *in vivo*.

Collectively, these results support a model whereby miRNA-mediated gene regulation, while dispensable to maintain normal colon homeostasis, becomes critical for its regeneration following acute damage.

### miRISC disruption impairs regeneration of the hematopoietic system

To further characterize the consequences of miRISC inhibition during tissue regeneration, we explored the possibility that other tissues may adopt a similar dynamic reliance on miRNA function.

Along with the intestinal epithelium, blood is one of the most rapidly turned over tissues in mice. Hematopoietic stem cells (HSCs) reside as a predominantly quiescent population in the bone marrow and are rapidly induced to re-enter the cell cycle in response to external cues, such as infection or injury^78^. Furthermore, HSCs can be readily isolated by flow cytometry and transplanted, allowing the study of mechanisms underlying regeneration at the single cell level.

To test the consequences of miRISC disruption in the regenerating hematopoietic system, we treated R26^T6B^ and R26^CTL^ mice on doxycycline-containing diet with a single dose of the cytotoxic drug 5-fluorouracil (5FU). 5-FU selectively depletes rapidly proliferating hematopoietic progenitors and leads to a compensatory increase in LT-HSC proliferation. Flow cytometry analysis of the bone marrow seven days after 5FU-injection showed that T6B expression prevented this compensatory increase in LT-HSC. We observed an identical phenotype when R26^T6B^ and R26^CTL^ mice that were bled repeatedly over a 3-week period to induce LT-HSC to re-enter the cell cycle (**Fig. 5a**).

**Figure 5.**
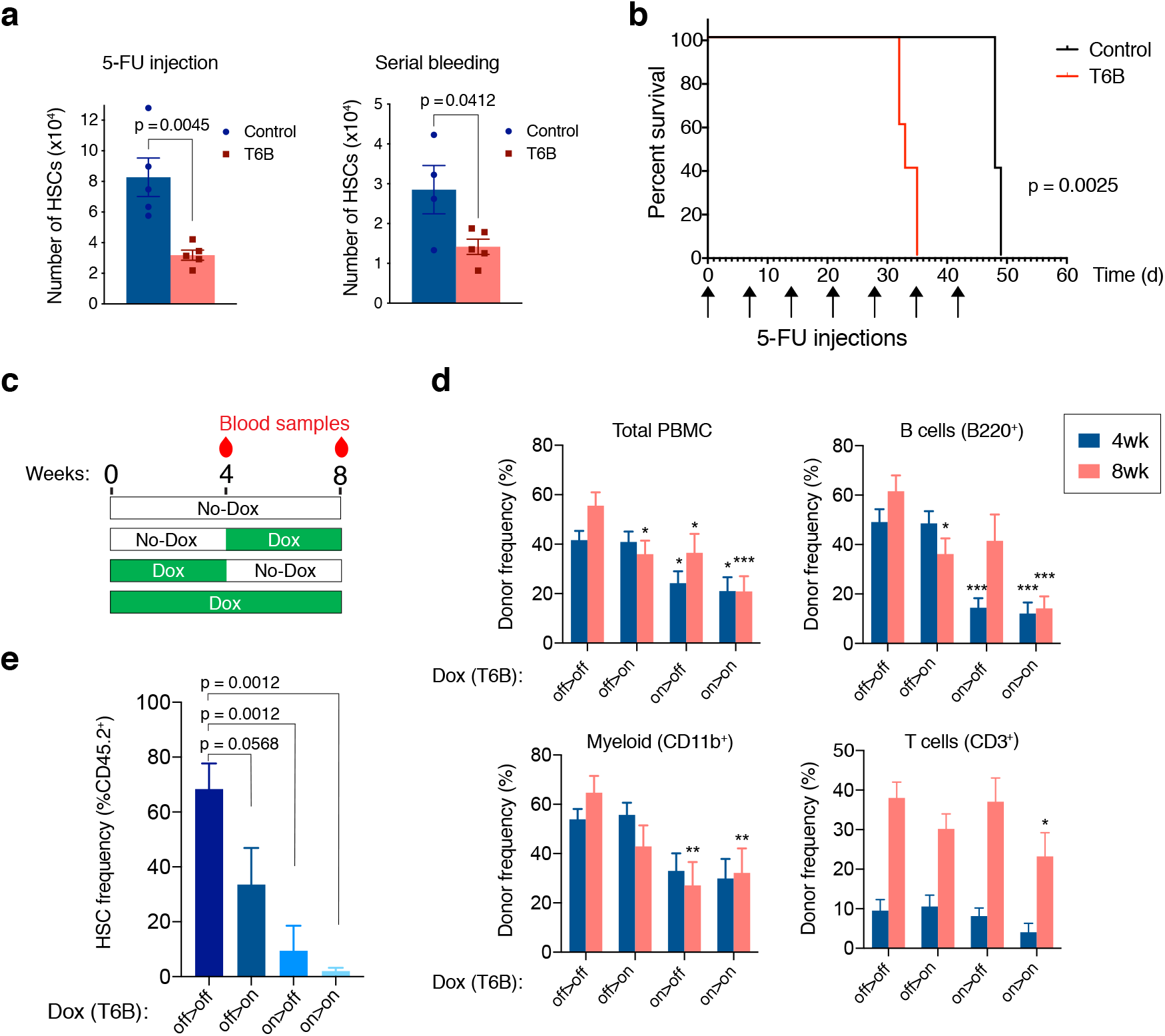
T6B-induced block of miRI SC assembly impairs the regeneration of the hematopoietic system. **(a)** Long term HSC in the bone marrow of R26^T6B^ and T6B^CTL^ mice treated with 5-FU or subjected to repeated bleeding (n = 5 for each genotype). Mice were maintained on doxycycline containing diet throughout the experiment. **(b)** Kaplan-Meier plots of R26^T6B^ (n = 5) and R26^CTL^ (n = 5) mice treated weekly with 5-FU for seven weeks. **(c)** Schematic of the bone marrow transplantation experiments: T6B was induced at different time points post-transplantation, and multi line age reconstitution was assessed at the indicated time points by FACS. **(d)** FACS analysis conducted on the peripheral blood of irradiated recipients transplanted 1:1 with T6B-expressing and wild-type bone marrow, and maintained on doxycycline diet according to scheme shown in panel c. Data are presented as mean ± s.d. *p < 0.05, **p < 0.01, ***p < 0.001, one-way ANOVA. off > off, n = 9; off > on, n = 10; on > off, n = 8; on>on, n = 8. **(e)** FACS analysis showing the frequency of T6B-extressing HSCs in the bone marrow of transplanted recipient mice kept on doxycycline diet according to scheme shown in panel c. off > off, n = 5; off > on, n = 5; on > off, n = 4; on > on, n = 5, one-way ANOVA.

The decreased number of HSCs in the bone marrow of R26^T6B^ mice after a single 5-FU challenge compared to controls, suggested that miRISC disruption impaired HSCs’ ability to re-enter the cell cycle and regenerate the hematopoietic compartment. Consistent with this hypothesis, when injected with repetitive 5-FU doses, R26^T6B^ mice showed significantly shorter survival compared to controls (**Fig. 5b**).

To more directly measure the regenerative capacity of HSCs in a context where T6B would only be expressed in hematopoietic cells, we performed competitive transplantation of T6B-expressing (CD45.2^+^) and wild-type (CD45.1^+^) bone marrows (1:1 ratio) into lethally irradiated hosts. The recipient animals were divided into four groups as shown in **Fig. 5c**: (i) a control group that was never administered doxycycline; (ii) a group maintained on a doxycycline-containing diet throughout the duration of the experiment (8 weeks); (iii) a group treated with doxycycline starting 4 weeks after transplant; and (iv) a group that was on doxycycline for only the first 4 weeks after transplant. Blood samples were taken at 4 and 8 weeks following the start of the experiment for analysis (**Fig. 5c**). This experiment was designed to test the prediction that expression of T6B during the first 4 weeks following transplant, when the regenerative demand is highest and when we hypothesize miRNA-mediated gene repression is required, would more severely affect the ability of donor cells to contribute to the recipient hematopoietic reconstitution compared to T6B expression after homeostasis is reestablished.

Consistent with this prediction, mice that were administered doxycycline in the first 4 weeks post-transplant had significantly fewer CD45.2^+^ peripheral blood mononuclear cells (PBMCs) (**Fig. 5d**). Contribution to the B cell population was particularly impaired by T6B expression but this was reversed once the recipients were taken off of doxycycline, consistent with the developmental block described earlier (**Fig. 3d** and **Extended Data Fig. 13**). Interestingly, the decrease in total CD45.2^+^ PBMCs and CD45.2^+^ myeloid cells was not reversed by doxycycline withdrawal, which suggested the T6B-expressing CD45.2^+^ HSCs might have been outcompeted by wild-type CD45.1^+^ HSCs in these recipients (**Fig. 5d**). Consistent with this hypothesis we observed a significant reduction in CD45.2^+^ HSCs only in the bone marrow of recipient animals that were fed a doxycycline-containing diet in the first 4 weeks post-transplant (**Fig. 5e**).

Taken together, these results support a model where the miRNA-mediated gene regulation is conditionally essential for the maintenance of hematopoietic stem cells during acute regeneration but is largely dispensable under homeostasis.

### An essential role for miRNA-mediated gene repression in the skeletal muscle and in the heart

As previously discussed, we observed low or no expression of T6B in the heart and skeletal muscle of R26^T6B^ mice treated with doxycycline (**Extended Data Fig. 4**), consistent with previous reports indicating that rtTA expression from the endogenous R26 promoter is tissue restricted^73^. To extend the analysis of the phenotype caused by the loss of miRISC activity to these tissues, we crossed T6B transgenic mice with the Rosa26-CAGs-rtTA3 strain^79^ in which the modified chicken beta-act-in with CMV-IE enhancer (CAG) promoter^80^ drives a more ubiquitous expression of the rtTA variant rtTA3 (hereafter CAG^T6B^). As expected, the pattern and intensity of T6B expression upon dox administration in CAG^T6B^ mice and R26^T6B^ mice were largely overlapping, except for the heart and the skeletal muscle, for which significant T6B expression was only observed in CAG^T6B^ mice (**Fig. 6a** and **Extended Data Fig. 4**). RNA-seq analyses confirmed inhibition of miRNA function in both heart and skeletal muscle of CAG^T6B^ mice upon dox administration (**Fig. 6b**).

**Figure 6.**
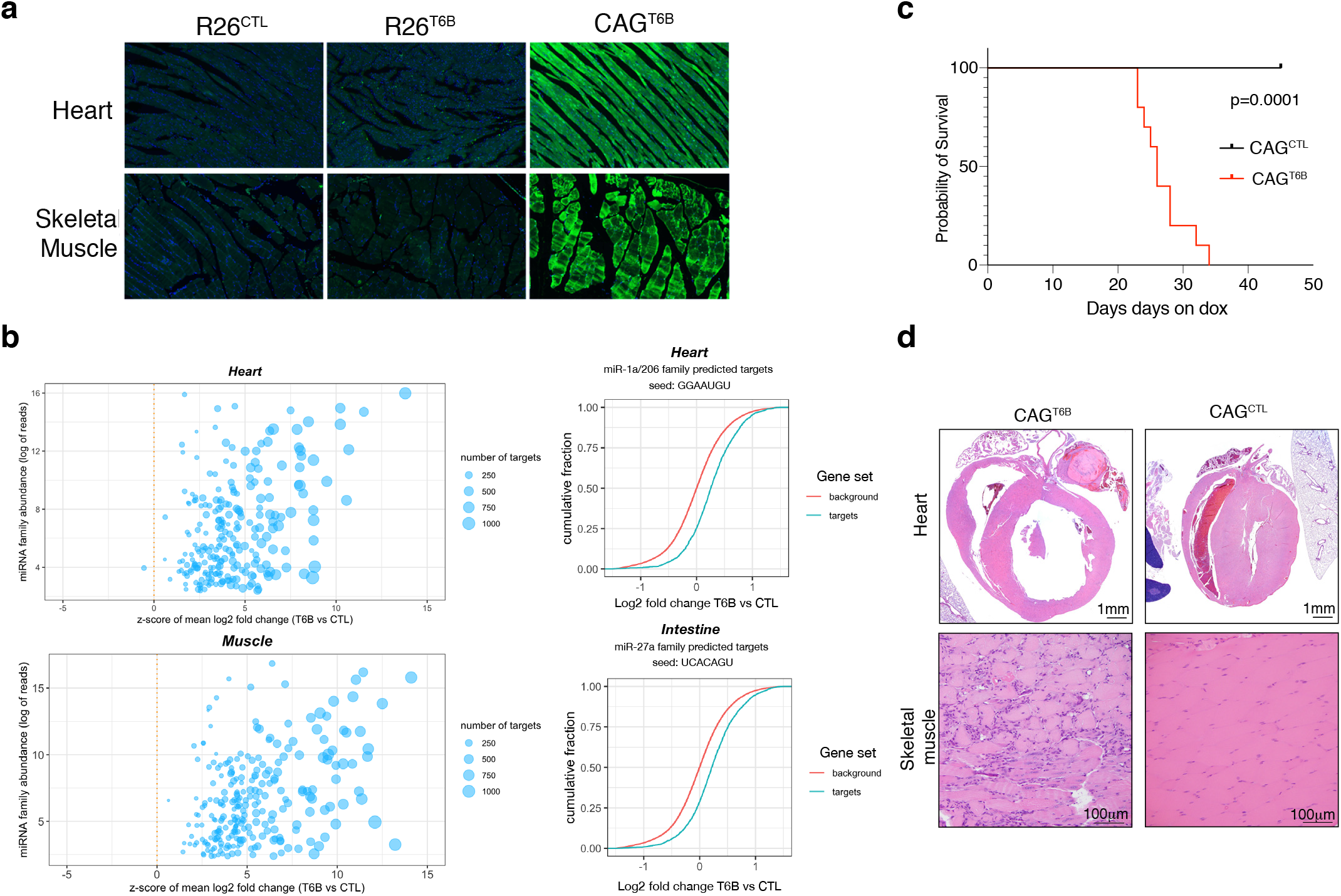
The miRNA pathway is essential in heart and skeletal muscle during homeostasis. **(a)** Detection of T6B expression with an anti-YFP antibody in the heart and skeletal muscle of R26^T6B^, CAG^T6B^, and R26^CTL^ mice maintained on doxycycline containing diet for 7 days. **(b)** Total RNA extracted from the heart (upper panel) and the skeletal muscle (lower panel) of CAG^CTL^ and CAG^T6B^ mice maintained on dox for 7 days was analyzed by RNAseq. Left panels: Scatter plot showing the effect of T6B expression on targets of conserved miRNA families were generated as described in figure 1d. The abundance of each miRNA family was calculated using dataset from Isakova et al.^100^. Right panels: Representative cumulative distribution plot of log2 fold changes in expression of predicted targets of the indicated miRNA families. **(c)** Kaplan-Meier curves of CAG^T6B^ and CAG^CTL^ mice (n = 8 for each genotype) maintained on doxycycline throughout the duration of the experiment. **(d)** Upper row: representative H&E staining showing marked dilation of the four cardiac chambers in hearts of CAG^T6B^ mice compared to controls (n = 9 for each genotype). Despite having thinner walls, the histomorphology of ventricular cardiomyofibers were within normal limits. Bottom row: representative H&E staining showing degenerative and regenerative changes in the skeletal muscle of the hind limbs of CAG^T6B^ mice compared to controls (n = 9 for each genotype).

In contrast to R26^T6B^ mice, CAG^T6B^ mice fed a doxycycline-containing diet showed a progressive decline in body mass (**Extended Data Fig. 16**), and died or reached a humane endpoint within 4-6 weeks (**Fig. 6c**). The decrease in body mass was not caused by intestinal malabsorption as, similarly to what observed in R26^T6B^ mice, we found no evidence of architectural defects throughout the intestine. In contrast, histopathologic examination of heart and skeletal muscle showed severe alterations in both organs, including dilated cardiomyopathy and diffuse muscular degeneration (**Fig. 6d**). All mice also showed necro-inflammatory changes in the liver, variable alterations in the pancreas, and increased urea nitrogen and alanine aminotransferase levels in the serum (data not shown). Such alterations are likely secondary to congestive heart failure, and/or to severe muscle catabolism as they were not observed in R26^T6B^ mice.

The emergence of severe cardiac and skeletal muscle phenotypes, as opposed to the lack of obvious structural and functional abnormalities in the majority of T6B-expressing tissues, points toward the existence of significant differences among adult tissues in their reliance on the miRNA pathway during homeostasis.

## Discussion

We report the generation of a novel genetically engineered mouse strain in which miRISC assembly and function can be temporally and spatially controlled in a reversible manner by a doxycycline-inducible transgene encoding a T6B-YFP fusion protein to address the role(s) miRNA-mediated gene regulation plays *in vivo* in adult tissues.

Surprisingly, in most adult tissues, we do not find an essential role for miRNA-mediated gene repression in organ homeostasis. A notable exception are the heart and the skeletal muscle, where miRISC inactivation in adult mice results in acute tissue degeneration and death even in the absence of tissue damage or exogenous stress.

Even though miRISC function is not overtly required for the homeostasis of other tissues, we have investigated the consequences of miRNA inhibition in the intestine and in the hematopoietic system of adult mice under homeostatic conditions and during tissue regeneration. These are tissues that periodically respond to external/internal stresses. In both tissues we have found that miRISC activity is dispensable for homeostasis. However, miRNA function becomes essential during tissue regeneration following acute injury. These results lend experimental support to the hypothesis that a major role for miRNA-mediated gene repression is to support tissue adaptation to stress.

In previous studies where *Dicer1* was conditionally ablated in the skeletal muscle of adult mice, muscle regeneration was impaired after acute injury, but no effect on muscle morphology or function was observed during homeostasis^24, 81, 82^. An explanation for this difference is that in the Dicer1 conditional knockout experiments miRNA levels were only partially reduced even weeks after *Dicer1* ablation, likely reflecting the high stability of these short non-coding RNAs. The T6B mouse strain we describe here overcomes this major limitation and allows the rapid and effective inhibition of miRNA-activity independently from the half-life of these molecules.

In this manuscript we have focused on the role of miRNA-mediated gene repression in adult mice. The same strategy for the acute inhibition of miRISC-activity can in principle be applied to other organisms. We have found that expression of T6B in embryos of both sea urchin (*Paracentrotus lividus*) and zebrafish (*Danio rerio*), induces developmental defects and gene expression changes consistent with the essential role of the miRNA pathway during development^83–89^ (**Extended Data Fig. 17**). Considering that *in vitro* T6B efficiently binds to AGO proteins from different non-mammalian organisms^58^, these findings are not unexpected, yet they highlight the usefulness of the T6B system for dissecting the miRNA pathway in a variety of animal models.

Despite its many advantages, the T6B mouse strain has also some unique limitations that need to be considered when designing and interpreting experiments.

First, although our biochemical and computational analysis of cells and tissues expressing T6B indicate that the peptide can effectively impair miRISC function, we cannot exclude some residual miRISC activity even in cells expressing high levels of the T6B transgene. The observation that we can recapitulate phenotypes observed in mice harboring complete targeted deletion of miR-143/145 miRNAs in the intestine^19^ and of miR-17~92 and miR-451 in the hematopoietic system^74, 75, 90^ is reassuring in this respect. For example, consistent with observations made in the regenerating intestine of miRNA-143/145 knockout mice^19^, we did not record any abnormalities or toxicity during the normal intestinal homeostasis of R26^T6B^ mice, whereas T6B expression became lethal during intestinal regeneration. Moreover, in the hematopoietic system, abnormalities were mostly restricted to B cell maturation, which are consistent with a developmental block at the Pro-B to Pre-B transition found in mir17~92 knockout mice^75^. Finally, we also observed a statistically significant decrease in hematocrit, erythrocyte volume and hemoglobin content in adult T6B-expressing mice, analogous to what reported in mice harboring targeted deletion of miR-451^74^.

In contrast, some of our results markedly differ from results obtained by conditional ablation of *Dicer1* in mice. For example, conditional knockout of *Dicer1* in the hematopoietic system has been reported to result in the rapid depletion of HSCs^91^. Furthermore, the lack of an overt phenotype in the intestine contrasts with previous reports showing that post-natal, conditional deletion of *Dicer1* results in depletion of Goblet cells^92, 93^, in addition to abnormal vacuolation and villous distortion in the small intestine^31, 93^. We cannot exclude that these differences are due to an incomplete inactivation of the miRISC pathway in T6B mice, but an alternative explanation is that they reflect the well-characterized miRNA-independent functions of DICER.

Another limitation to be considered is the possibility that T6B expression impairs the activity of other complexes in addition to the miRISC. Although RNAseq analysis of cells expressing T6B has not revealed changes that are not explained by loss of miRNA-mediated gene repression and the phenotypes observed are consistent with loss of miRNA activity, this possibility cannot be formally excluded at this time. Further studies to experimentally identify T6B interactors in cells and tissues will be important to formally address this possibility.

In conclusion, we have developed a novel mouse strain that enables investigating the role of miRNA-mediated gene repression in adult organisms. The body of data presented here suggest that in adult animals miRNAs primarily provide for the ability to adaptively change gene expression in response to the physiologic and pathologic stresses that accompany metazoans’ life. It is likely that the specific miRNAs and stresses differ based on the adult organ or tissue being studied and the model we have generated will be useful address these important aspects of miRNA biology.

## Methods

### Animal models

The Rosa26^rtTA/rtTA^; ColA1^T6B/T6B^ (R26^T6B^) mice were generated by site-specific integration of the transgene coding for the FLAG-HA-T6B-YFP fusion protein within the Col1a locus of KH2 embryonic stem cells (Col1A-frt/ Rosa26 rtTA)^72^. Briefly, the FLAG-HA-T6B-YFP (FH-T6B-YFP) DNA fragment was subcloned into the targeting vector, as described in “Vectors and molecular cloning”. A mixture of 5μg of the targeting vector and 2.5μg of the pCAGGS-flpE-puro (Addgene #20733), Flippase recombinase-expressing vector were electroporated into KH2 cells, using 4D-Nucleofector core unit (Lonza), following manufacturer’s “Primary cells P3” protocol. Selection of targeted clones was initiated 48h after electroporation, using 150μg hygromycin per mL of culture medium. 10 days later, individual hygromycin-resistant ES cell clones were analyzed by PCR to confirm correct integration of the knock-in allele. Clones carrying the correctly integrated knock-in allele were genotyped using a three-primer PCR, with the following primers: 1) 5’-AATCATCCCAGGTGCACAGCATTGCGG-3’; 2) 5’-CTTTGAGGGCTCATGAACCTCCCAGG-3’; 3) 5’-ATCAAGGAAAC-CCTGGACTACTGCG-3’ A 287pb-long PCR product indicates successful integration of the transgene into the Cola1, while a 238bp-long PCR product indicates a wild type, untargeted locus. Two independent ES clones were injected into C57BL/6J albino blastocysts and backcrossed the resulting chimeras to C57BL/6J mice to achieve germline transmission of the recombinant allele. F1 animals were then intercrossed to generate animals expressing rtTA from the R26 locus under control of the R26 endogenous promoter, while expressing the T6B fusion protein from the Col1a locus under control of the tetracycline-responsive element (TRE) and the minimal CMV promoter. Animals were genotyped as follows: to assess the presence of the transgene in the ColA1 locus, PCR was carried out as for the genotyping of KH2 cells. To assess the presence of the rtTA transgene in the Rosa26 locus, a three-primer PCR was performed, using the following primers: 1) 5’-AAAGTCGCTCTGAGTTGTTAT-3’; 2) 5’-GCGAA-GAGTTTGTCCTCAACC-3’; 3) 5’-CCTCCAATTTTACACCTGTTC-3’ A 350pb-long PCR product indicates the presence of the rtTA transgene into the Rosa26 locus, while a 297bp-long PCR product indicates the presence of a wild type locus. CAG^rtTA/rtTA^; Col1A^T6B/T6B^ (CAG^T6B^) mice were generated by backcrossing R26^T6B^ with Rosa26-CAGs-rtTA3 mice (a gift from Scott Lowe, MSKCC). In the Rosa26-CAGs-rtTA3 mice, the knock-in allele has the CAG promoter driving the expression of the third-generation reverse tetracycline-regulated transactivator gene (rtTA3), all inserted into the Gt(ROSA)26Sor locus. *In vivo* doxycycline-dependent expression of the FLAG-HA-T6B-YFP transgene was achieved by feeding mice chow that contained doxycycline at the concentration of 625mg/Kg (Envigo #TD01306). Mice were maintained and euthanized in accordance with a protocol approved by the Memorial Sloan-Kettering Cancer Center Institutional Animal Care and Use Committee.

### Necropsy, staining and histopathology

Mice were euthanized with CO2. Following gross examination all organs were fixed in 10% neutral buffered formalin, followed by decalcification of bone in a formic acid solution (Surgipath Decalcifier I, Leica Biosystems). Tissues were then processed in ethanol and xylene and embedded in paraffin in a Leica ASP6025 tissue processor. Paraffin blocks were sectioned at 5 microns, stained with hematoxylin and eosin (H&E), and examined by a board-certified veterinary pathologist. The following tissues were processed and examined: heart, thymus, lungs, liver, gallbladder, kidneys, pancreas, stomach, duodenum, jejunum, ileum, cecum, colon, lymph nodes (submandibular, mesenteric), salivary glands, skin (trunk and head), urinary bladder, uterus, cervix, vagina, ovaries, oviducts, adrenal glands, spleen, thyroid gland, esophagus, trachea, spinal cord, vertebrae, sternum, femur, tibia, stifle join, skeletal muscle, nerves, skull, nasal cavity, oral cavity, teeth, ears, eyes, pituitary gland, brain. To detect goblet cells in the intestine, the AB/PAS kit (ThermoFisher #87023) was used according to the manufacturer’s instructions.

### Immunofluorescence

For the staining of intestine sections shown in Figure 3 and Extended Data Figure 8, formalin-fixed, paraffin-embedded (FFPE) slides were deparaffinized and rehydrated according to a standard xylene/ethanol series. After heat-induced epitope retrieval in sodium citrate (pH6), tissue sections were permeabilized in triton X-100, blocked, and incubated with the following 1° antibodies: PH3 (Cell Signaling #970) at 1:200 dilution; Lysozyme (ThermoFisher #RB-372-A1) at 1:200 dilution; E-Cadherin (BD#610181) at 1:750 dilution; YFP (Invitrogen #A11122) at 1:250 dilution; Ki67 (Cell Signaling #12202) at 1:400 dilution. Next, cells were washed with PBS containing 0.05% Triton X, and incubated with the following 2° antibodies: Goat anti-Rabbit IgG, Alexa Fluor 488 (ThermoFisher #A11034) at 1:250 dilution; Goat anti-Mouse IgG2a, Alexa Fluor 594 (ThermoFisher #A11029) at 1:250 dilution. For the staining of tissue sections shown in Figure 2, Figure 4 and Extended Data Figure 4, FFPE tissue sections were cut at 5 μm and heated at 58°C for 1 hr. The antibody against GFP (Abcam, ab13970, 2μg/ml) was incubated for 1 hr and detected with Leica Bond RX. Appropriate species-matched secondary antibody and Leica Bond Polymer anti-rabbit HRP were used, followed by Alexa Fluor 488 tyramide signal amplification reagent (Life Technologies, B40953). After staining, slides were washed in PBS and incubated in 5 μg/ml 4’6-diamidino-2-phenylindole (DAPI) (Sigma Aldrich) in PBS (Sigma Aldrich) for 5 min, rinsed in PBS, and mounted in Mowiol 4–88 (Calbiochem). Slides were kept overnight at −20°C before imaging.

### Immunohistochemistry

For IHC, deparaffinized sections were subjected to antigen retrieval and processed with the EnVision+ HRP kit (K401111–2, DAKO, Glostrup, Denmark) according to the manufacturer’s instructions. A primary polyclonal antibody against Ki67 (Cell Signaling #12202) at 1:400 dilution was diluted in Antibody Diluent (DAKO #S0809) and incubated overnight at 4°C. Next, sections were incubated in the provided anti-rabbit HRP-labeled polymer reagent and detection was performed according to the manufacturer’s protocol. Images were acquired using an Olympus BX-UCB slide scanner.

### *In situ* hybridization

5μm sections were obtained from formalin-fixed, paraffin-embedded (FFPE) colons from age/sex-matched mice. Before staining, tissue slides were deparaffinized, rehydrated and permeabilized according to standard procedures. Detection was carried out using RNAscope 2.5 HD Detection Reagent, BROWN (ACD # 320771), with a specific RNAScope Igfbp5 Probe (ACD #425738, according to the manufacturer’s instructions.

### Serum chemistry and hematology

For serum chemistry, blood was collected into tubes containing a serum separator, the tubes were centrifuged, and the serum was obtained for analysis. Serum chemistry was performed on a Beckman Coulter AU680 analyzer and the concentration of the following analytes was determined: alkaline phosphatase, alanine aminotransferase, aspartate aminotransferase, creatine kinase, gamma-glutamyl transpeptidase, albumin, total protein, globulin, total bilirubin, blood urea nitrogen, creatinine, cholesterol, triglycerides, glucose, calcium, phosphorus, chloride, potassium, and sodium. Na/K ratio, albumin/globulin ratio were calculated. For hematology, blood was collected retro-orbitally into EDTA microtainers. Automated analysis was performed on an IDEXX Procyte DX hematology analyzer.

### Dextran sulfate sodium (DSS) treatment and post DSS treatment quantitative analyses

Mice kept in doxycycline-containing chow were treated for 5 days with 4% w/v DSS (FW 40.000) (Cayman Chemical #23250) dissolved in drinking water. Body mass was monitored daily. Measurements of colon length, aggregated length of ulcers, percentage of colon with ulcers, area of ulcers, the number of immune nodules and the area of immune nodules were obtained using OMERO (https://www.openmicroscopy.org/omero/). Measurements of these parameters were used to estimate the extent of damage and colitis induced by DSS treatment. All measurements were acquired from H&E-stained colon sections. Ulcer was defined as regions of colon with complete/partial loss of formal epithelial structural, accompanied by massive immune infiltrates. Colon length was measured by tracing the length of muscular layer of each colon. Length of ulcer was measured as the added length of each ulcerated region along the colon. Ulcer percentage was calculated as the length of ulcer/length of colon. The area of each individual ulcer was also measured and summed for each animal. Clear immune nodules are visible, showing aggregates of immune cells with high nucleus/cytoplasm ratio. Number and area of the immune nodules were summarized for each animal.

### Tissue isolation and total lysates preparation

Organs extracted from 8- to 12-week-old mice, perfused with PBS, were snap-frozen in liquid nitrogen and stored at −80 °C until further processing. To prepare total extract from solid tissues, tissues were pulverized using a mortar, resuspended in 1mL of lysis buffer per cm3 of tissue, and dounce-homogenized with a tight pestle until completely homogenized. Next, extracts were cleared by centrifugation at 20,000 × g for 5 min followed by a second step of centrifugation at 20,000 × g for 5 min. To prepare total extracts from cultured cells, pelleted cells were snap frozen in liquid nitrogen and stored at −80 °C until further processing. Pellets were then resuspended in lysis buffer, incubated for 10 minutes on ice, and cleared by centrifugation at 20,000 × g. Two different lysis buffers were used, depending on the specific downstream application. For IP and size exclusion chromatography, lysates were prepared in SEC buffer (150 mM NaCl, 10 mM Tris-HCl pH 7.5, 2.5 mM MgCl2, 0.01% Triton X-100). For Western blotting applications, lysates were prepared in RIPA buffer (Sigma-Aldrich # R0278). Upon usage, both buffers were supplemented with the addition of EDTA-free complete protease inhibitors (Sigma-Aldrich #11836170001), phosphate inhibitors (Roche #04906837001), and 1mM DTT.

### Cell Lines and Culture Conditions

Cell lines were maintained in log-phase growth in a humidified incubator at 37°C, 5% CO2 prior to experimental manipulation. HCT116 colorectal adenocarcinoma cells were maintained in McCoy’s medium supplemented with 10% heat-inactivated fetal calf serum (FCS, GIBCO, Cat#16141079), 10 U/ml penicillin/streptomycin, and 2 mM L-glutamine. Mouse embryonic fibroblasts (MEF) were grown in Dulbecco’s Modified Eagle Medium (DMEM) supplemented with 10% heat-inactivated fetal calf serum (FCS, GIBCO), 10 U/ml penicillin/streptomycin, and 2 mM L-glutamine. KH2 embryonic stem cells were cultured in gelatin-coated plates in presence of irradiated DR4 Mouse Embryonic Fibroblasts (ThermoFisher #A34966), and maintained in Knock-Out DMEM (GIBCO, Cat#10829018), supplemented with 15% FCS (GIBCO), GlutaMax (GIBCO Cat#35050061), 100 μM non-essential amino acids (Sigma-Aldrich Cat#M7145), 1000 U/mL leukemia inhibitory factor (LIF, Millipore Cat#ESG1107), 10U/mL penicillin/streptomycin (GIBCO Cat#15070063) and 100 mM 2-Mercaptoethanol (Bio-Rad Cat#1610710), and nucleosides (Millipore Cat#ES-008-D).

### Flow cytometry

Analysis of bone marrow populations was performed by harvesting femurs and tibiae from euthanized mice. Bone marrow was isolated by centrifugation (REF), resuspended in FACS buffer (PBS with 2% fetal calf serum) and passed through a 40μm cell strainer to make a single cell suspension. Nonspecific antibody binding was blocked by incubation with 10μg/ml Rat IgG (Sigma #I-8015) for 15 min on ice. Antibodies used to identify HSCs included a cocktail of biotinylated lineage antibodies (Gr1, CD11b, TER119, B220, CD3, CD4, CD8), CD117 (c-kit) APC (2B8), Sca-1 (D7) PE-cy7, CD150 PE, and CD48 Pacific Blue. B cell progenitors were identified with the following antibodies: B220, CD19, CD25, CD43, IgM, IgD and c-kit. For analysis of peripheral blood mononuclear cells, blood was collected retro-orbitally from live mice into EDTA microtainers. Whole blood was lysed in ACK buffer for 5 min at room temperature, washed with FACS buffer and pelleted prior to antibody staining. Mature blood populations were identified with the following antibodies: CD45.1, CD45.2, Gr1, CD11b, B220, CD3. Cells were incubated with primary antibodies for 45 min, washed once with FACS buffer and incubated with BV711 streptavidin conjugate for 15 min. All incubations were carried out on ice and protected from light. Antibodies were purchased from Biolegend or eBioscience.

### Bone marrow transplantation

8–12-week-old CD45.1^+^ C57BL/6 (BoyJ) mice (JAX) were lethally irradiated by exposure to 1100cGy of gamma irradiation from a cesium source, administered in two doses, split 4h apart. Bone marrow suspensions from CAG^T6B^ (CD45.2^+^) and BoyJ mice were counted, mixed 1:1 and transferred intravenously by retro-orbital injection into isofluorane-anesthetized, irradiated recipients.

### Size exclusion chromatography (SEC)

SEC was performed using a Superose 6 10/300 GL prepacked column (GE Healthcare) equilibrated with SEC buffer. A total of 400μL (1.5–2 mg) of precleaned total extracts either from cultured cells or tissues were run on the SEC column at a flow rate of 0.3 mL/min. 1mL fractions were collected. Proteins were extracted from each fraction by TCA precipitation following standard procedures, and run on SDS-PAGE gels for Western blotting analysis.

### Western blotting and antibodies

Western blotting was performed using the Novex NuPAGE SDS/PAGE gel system (Invitrogen). Total cell lysates were run either on 3–8% Tris-acetate or 4–12% Bis-Tris precast gels, transferred to nitrocellulose membranes, and probed with antibodies specific to proteins of interest. Detection and quantification of blots were performed on Amersham hyperfilm ECL (Cytiva #28906839) and developed on film processor SRX-101A (Konica). Antibodies used for Western blots were obtained from commercial sources as follows: anti-GW182 (Bethyl #A302-239A), anti-Ago2 (Cell Signaling #2897), anti-PABP1 (Cell Signaling #4992), anti-RPL26 (Bethyl #A300-686A), anti-GAPDH (Sigma #G8795), anti-βActin (Sigma #A2228) anti-GFP (Roche #11814460001), anti-Tubulin (Sigma-Aldrich #T9026) anti-HA (Cell Signaling #C29F4), anti-Rabbit IgG, HRP-conjugated (GE Healthcare #NA934), anti-Mouse IgG, HRP-conjugated (GE Healthcare #NA931).

### Immunoprecipitation (IP)

For IP of AGO-T6B complexes from human HCT116 cells, 500μg of lysates in 500 μL of SEC buffer were incubated for 3 hours with primary antibodies directed to either AGO proteins (WAKO anti-AGO2 #011-22033, EMD Millipore anti-panAGO #MABE56) or directed to T6B-fusion protein (Cell Signaling anti-FLAG #8146S, Cell Signaling anti-HA #2367S) or mouse IgG1 isotype control (Cell Signaling #5415). Next, lysates were incubated with 20μl of protein A/G PLUS-Agarose beads (Santa Cruz #2003) for 1 hour. For IP of AGO-T6B complexes from mouse tissues, 500μg of lysates in 500 μL of SEC buffer were incubated for 2 hours with GFP-trap magnetic agarose beads (Chromotek #gtma-10) or binding control beads (Chromotek #bmab-20). The immune complexes were run on SDS-PAGE and analyzed by Western blotting.

### Vectors and molecular cloning

The targeting vector expressing the FH-T6B-YFP under control of TRE and CMV minimal promoter, was generated from a modified version of the pgk-ATG-frt plasmid (Addgene plasmid #20734), in which the region of pgk-ATG-frt comprised between the EcoRI site and the PciI site was substituted with the rabbit β-globin polyadenylation signal (RBG pA). The FH-T6B-YFP DNA insert was generated by PCR using the plasmid pIRES-Neo-FH-T6B-YFP^58^ as a template. PCR was carried out using the following primers: Forward: 5’-GACTACAAGGACGACGATGACAAG-3’, Reverse: GTTACTTG-TACAGCTCGTCCATG. Next, the modified pgk-ATG-frt, was cut with NcoI, filled-in to produce blunt ends, dephosphorylated and ligated to the PCR-generated FH-T6B-YFP DNA fragment, according to standard subcloning procedures. Below, a scheme of the cloning strategy:

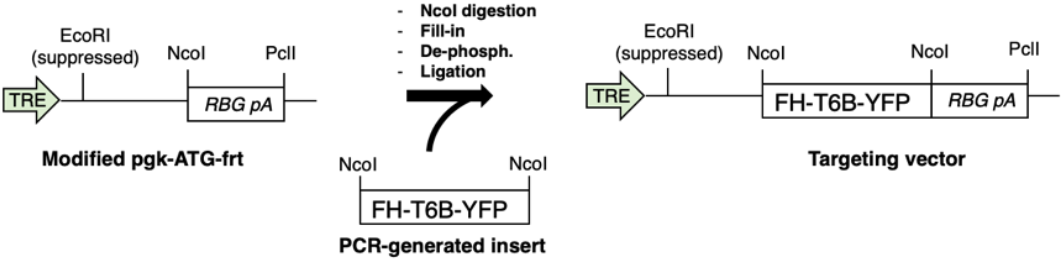

To generated cell lines expressing either FH-T6B-YFP or FH-T6B^Mut^-YFP fusion proteins in a doxycycline-inducible manner, a modified version of the retroviral vector pSIN-TREtight-HA-UbiC-rtTA3-IRES-Hygro (hereafter TURN vector, a gift from Scott Lowe) was used to transduce commercially available HCT116 and MEFs cell lines. TURN is an all-in-one Tet-on vector that includes: 1) The rtTA3 gene under the human ubiquitin C promoter; 2) The transgene of interest driven by a tetracycline-responsive element (TRE)/CMV promoter. We used the pIRES-Neo-FH-T6B-YFP described in Hauptmann ae al.^58^ as a template to generate by PCR the DNA fragments coding either for FH-T6B-YFP or for FH-T6BMut-YFP fusion proteins. DNA fragments were then inserted into the XhoI/EcoRI-digested TURN vector to generate TURN^T6B^ and TURN^T6Bmut^ vectors used for the transduction of parental HCT116 and MEFs.

### Small RNA Transfection

Silencer GAPDH siRNA (ThermoFisher AM4624) and Silencer Select Negative Control 1 siRNA (ThermoFisher AM4611). Small RNAs were transfected at 10 pM per 1 × 10^6^ cells. MEFs were reverse transfected using Lipofectamine RNAiMAX. Lipofectamine RNAiMAX was combined with 20 μM small RNAs at a 4:3 ratio (vol:vol) in Opti-MEM and incubated for 20 min at room temperature. Trypsinized cells were added to culture dishes containing siRNAs and Lipofectamine RNAiMAX at 3.8 × 10^4^ cells per centimeter squared. Three volumes of complete medium were added to culture dishes and cells were incubated for 2–3 days before further processing.

### Small RNA sequencing

Total RNA was extracted from T6B and induced as well uninduced T6B^mut^ MEFs and 1 μg was used as input for sRNA-seq library preparation as described in ref. ^94^. Briefly, 1 μg total RNA was ligated to nine distinct pre-adenylated 26-nt 3’-adapters with a 5-nt barcode using a mutated and truncated Rnl2 followed by urea gel purification and size selection and 5’-adapter ligation with Rnl1. This ligation reaction was again gel purified and size-selected for fully ligated product and reverse transcribed using SuperScript III RT followed by PCR amplification using Taq polymerase for 25 cycles. The final PCR product was separated on a 2% agarose gel in TBE buffer and extracted using the QIAgen gel extraction kit according to the manufacturer’s instructions including all optional steps. After high-throughput sequencing, small RNA reads were aligned to a miRNA genome index built from 1,915 murine pre-miRNA sequences from miRbase version 21^95^ (ftp://mirbase.org/pub/mirbase/21/) using Bowtie v2.4.296. Mature miRNA abundance was calculated by counting reads falling within 4 bps at each of the 5’ and 3’ end of the annotated mature miRNAs. miRNA seed family data were downloaded from the TargetScan website at http://www.targetscan.org/mmu_71/mmu_71_data_download/miR_Family_Info.txt.zip. For miRNA family level analysis, read counts mapping to members of the same miRNA family were summed up.

### RNAseq analysis

Total RNA from heart, skeletal muscle, colon and liver of sex-matched littermate animals, and total RNA from cell lines was extracted using TRIzol Reagent (Invitrogen) according to manufacturer’s instructions and subjected to DNase (QIAGEN) treatment. After RiboGreen quantification and quality control by Agilent BioAnalyzer, 500ng of total RNA with RIN values of 7.0-10 underwent polyA selection and TruSeq library preparation according to instructions provided by Illumina (TruSeq Stranded mRNA LT Kit, catalog # RS-122-2102), with 8 cycles of PCR. Samples were barcoded and run on a HiSeq 4000 in a PE50/50 run, using the HiSeq 3000/4000 SBS Kit (Illumina). An average of 34 million paired reads was generated per sample. The percent of mRNA bases averaged 60% over all samples. Reads were aligned to the standard mouse genome (mm10) using STAR v2.5.3a^97^. RNA reads aligned were counted at each gene locus. Expressed genes were subjected to differential gene expression analysis by DESeq2 v1.20.0 ^98^. Gene expression analysis was performed by comparing tissues from T6B-expressing animals (R26^T6B^ or CAG^T6B^) with relative littermate controls.

### Z-score calculation

For conserved miRNA families, the mean log2-fold change of predicted targets compared to the rest of the transcriptome (back-ground) was calculated. The means were converted to z-scores using an approach developed by Kim and Volsky^99^. Z-score = (Sm -m)3m1/2/ SD, where Sm is the mean of log2-fold changes of genes for a given gene set, m is the size of the gene set, and mand SD are the mean and the standard deviation of background log2-fold change values.

## Acknowledgements

This work was funded by the Starr Foundation’s Tri-Institutional Stem Cell Initiative (A.V., T.L., D.B. and T.T.), and by the NIH/NCI (grants R01CA149707 and R01CA245507 to A.V. and P30 CA008748 to C.B.T.). Y.M. was supported by a Medical Scientist Training Program grant from the National Institute of General Medical Sciences of the National Institutes of Health under award number: T32GM007739 to the Weill Cornell/Rockefeller/Sloan Kettering Tri-Institutional MD-PhD Program. We acknowledge the use of the following core facilities at the Memorial Sloan Kettering Cancer Center (MSKCC): The Molecular Cytology Core; The Mouse Genetics Core Facility; The Laboratory of Comparative Pathology, and the Integrated Genomics Operation Core, funded by the NCI Cancer Center Support Grant (CCSG, P30 CA08748), Cycle for Survival, and the Marie-Josée and Henry R. Kravis Center for Molecular Oncology. We thank Sebastien Monette and Ileana Miranda for their contribution in the phenotypic analysis of R26^T6B^ and CAG^T6B^ mouse strains; Davide Pradella, Rui Gao, Saurabh Yadav and members of the Benezra laboratory for discussion and suggestions.

## Author contribution

G.L.R., B.K. and A.V. designed the research project; Methodology: G.L.R., B.K., T.T., G.M., C.B.T., T.L. and A.V.; Investigation: G.L.R., B.K., B.S., X.L., M.Z., K.A., V.C., Y.M., V.A. and J.A.V.; Formal Analysis and Software: X.L., B.S., D.B. and A.V.; Writing – Original Draft: G.L.R., B.K. and A.V.; Writing – Review & Editing: G.L.R., B.K., B.S., X.L., M.Z., K.A., V.C., D.B., V.A., J.A.V., T.T., G.M., C.B.T., T.L., K.M.H., and A.V.; Funding Acquisition: C.B.T., T.T., T.L., D.B. and A.V.; Resources: P.O., K.C. and C.M.; Supervision: K.M.H. and A.V.

## Supplementary information

**Extended Data Fig. 1.**
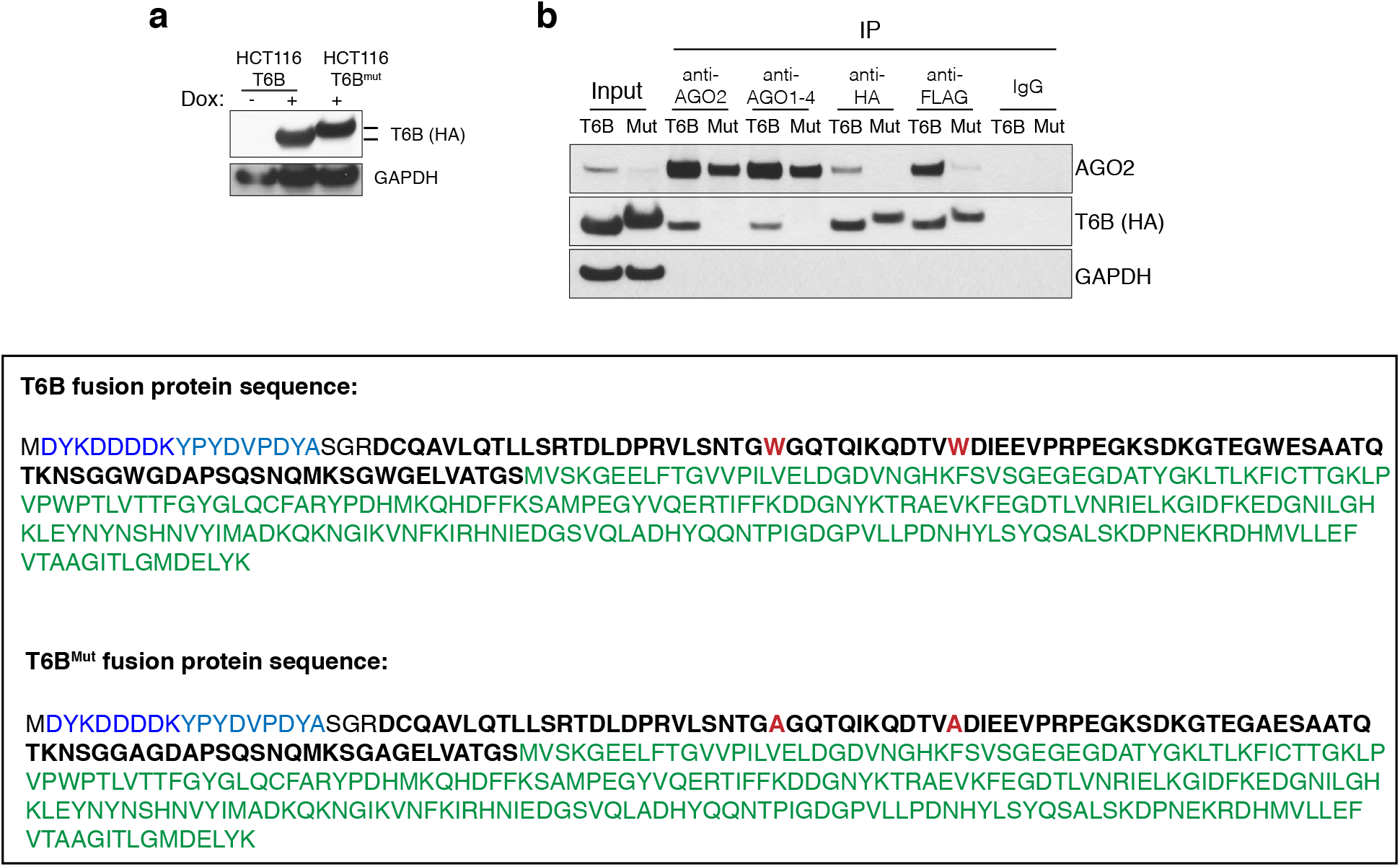
**(a)** HCTT116 cells transduced with retroviral vectors expressing a doxycycline-inducible T6B or T6B^Mut^ transgene (FH-T6B-YFP) were cultured in the presence of doxycycline for 48. Whole cells lysates were probed with an anti-HA antibody. **(b)** Lysates from (a) were immunoprecipitated with the indicated antibodies and blotted against AGO2, FH-T6B-YFP (anti-HA) and GAPDH. Note that the T6B fusion protein, but not its mutant version (T6B^Mut^), binds to AGO proteins. Lower panel. Aminoacid sequence of the T6B and T6B^Mut^ fusion proteins. Both T6B versions have HA and FLAG tags at the N termini, and are fused to the yellow fluorescent protein (YFP) at the C-termini. In T6B^Mut^, all Tryptophan residues (red) are mutated to Alanine to prevent interaction with AGO proteins. Blue, FLAG-tag; light blue, HA-tag; bold black, T6B; green, YFP.

**Extended Data Fig. 2.**
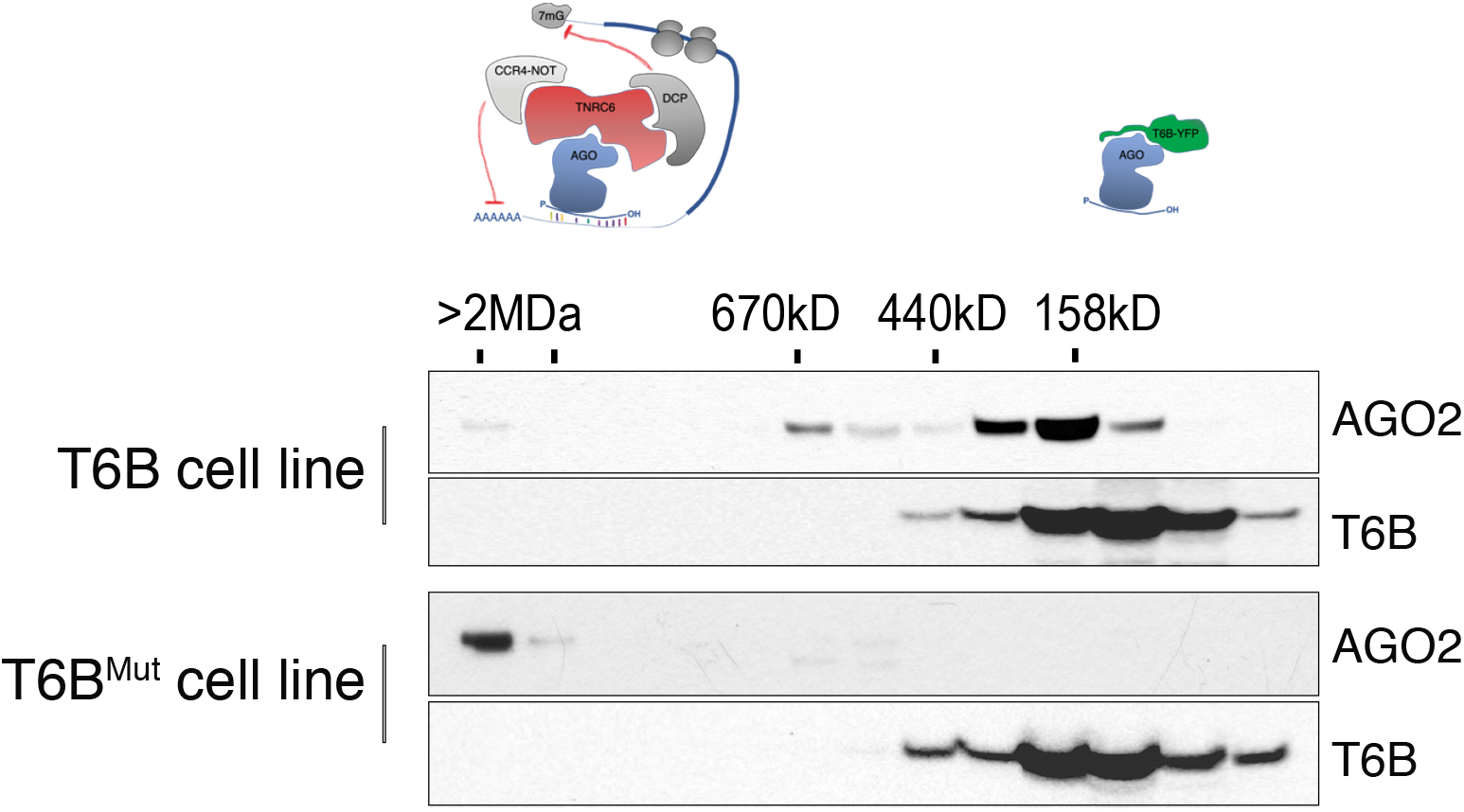
Size exclusion chromatography was performed on whole cell lysates from MEFs transduced with retroviral vectors expressing a doxycycline-inducible T6B or T6B^Mut^ transgene and cultured in presence of doxycycline for 48h. Eluted fractions were probed with the anti-AGO2 or anti-HA antibodies to determine the elution profile of AGO2 and T6B, respectively.

**Extended Data Fig. 3.**
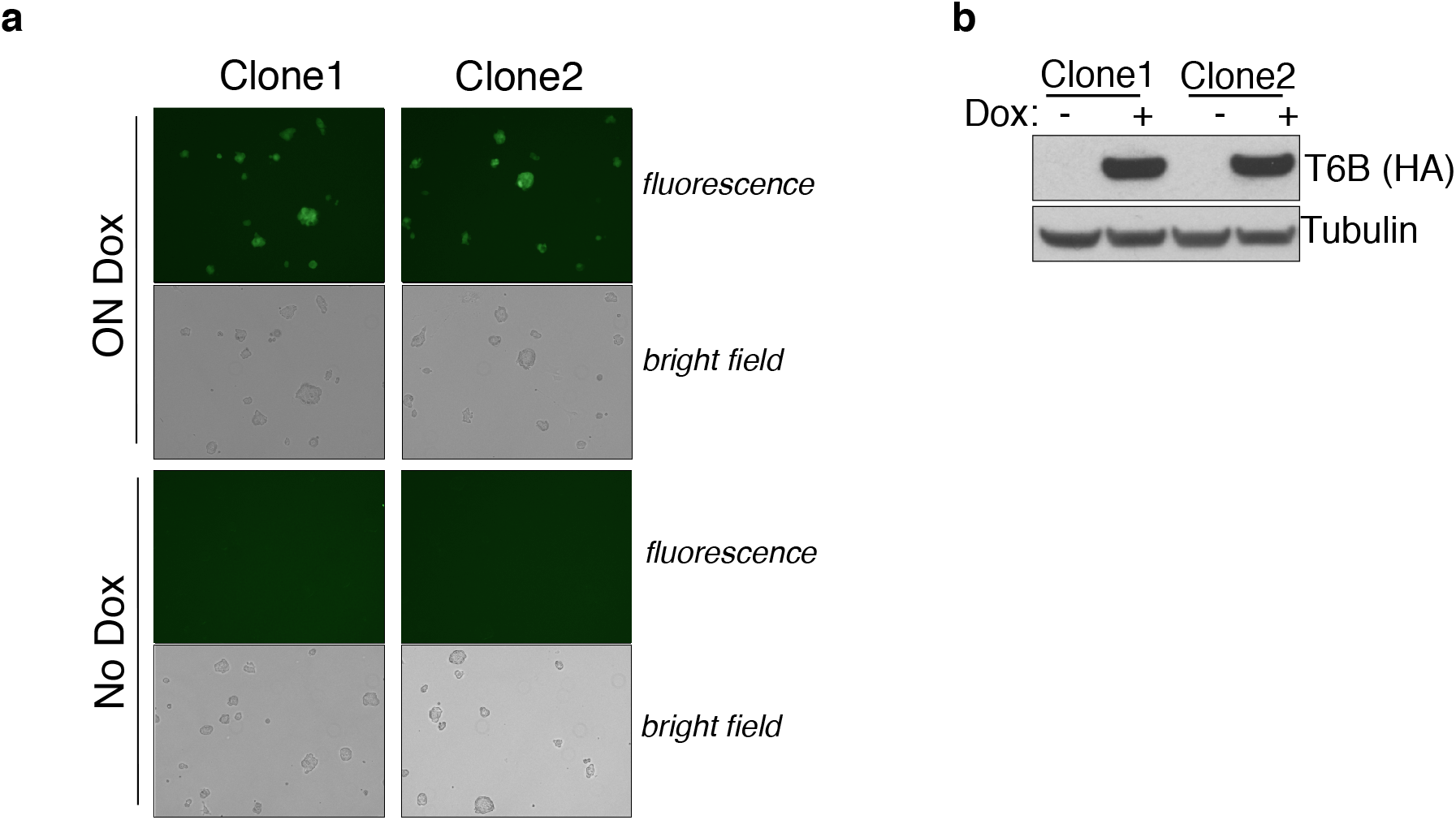
**(a)** Two independent targeted ES clones were cultured in the presence or absence of doxycycline for 48h and examined by epifluorescence microscopy to detect FH-T6B-YFP expression. The same exposure was used for all images. Bright field images are also shown for each clone. **(b)** Whole cell lysates from the clones shown in (a) were reprobed with an anti-HA antibody to detect expression of the T6B fusion protein.

**Extended Data Fig. 4.**
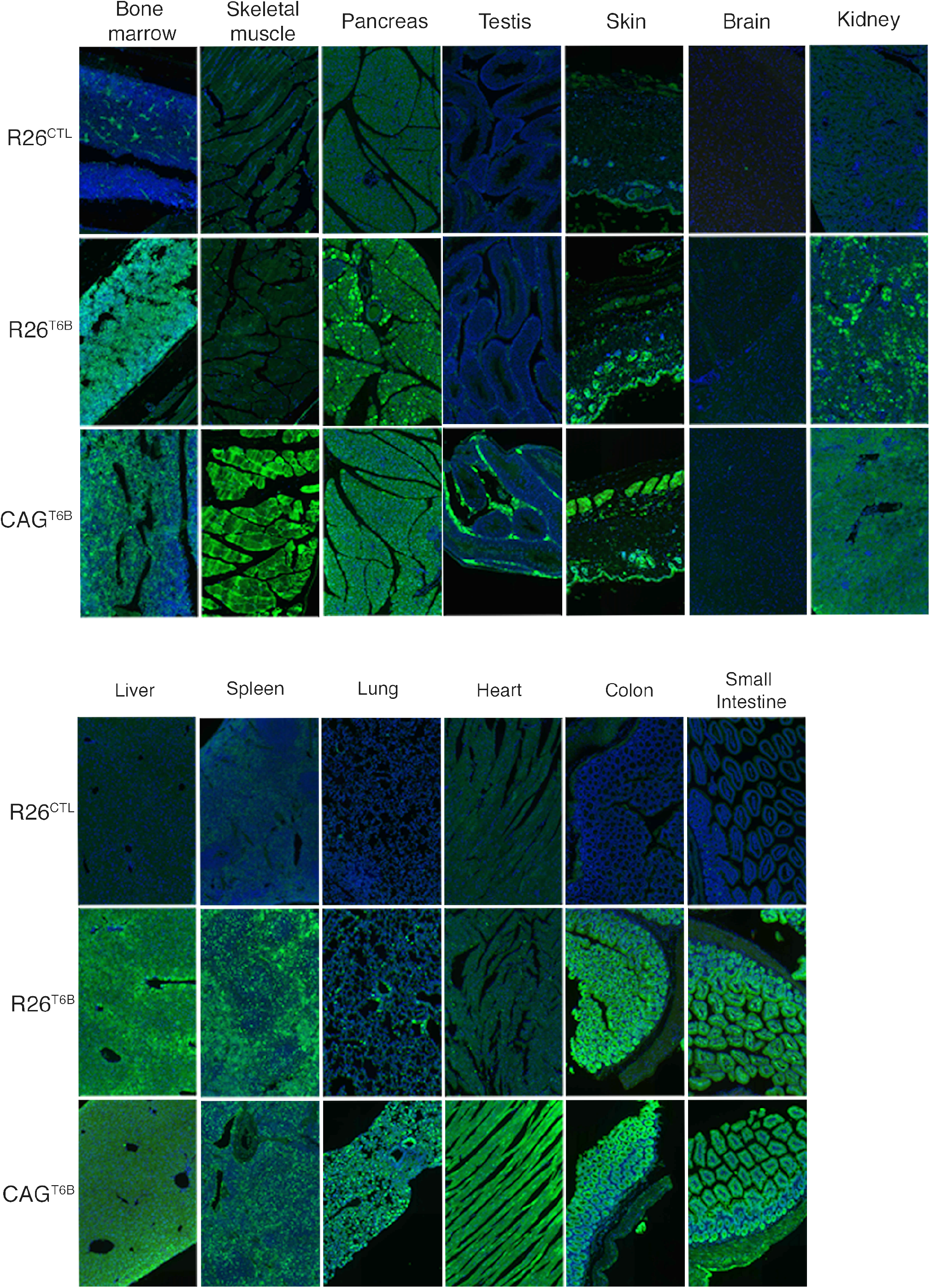
Immunofluorescence imaging using a YFP-specific antibody, showing T6B expression in a panel of tissues of adult R26^T6B^ mice (second column) and CAG^T6B^ mice (third column) fed doxycycline-containing diet for 7 days. Tissues from R26^CTL^ (first column) mice fed doxycycline-containing diet for 7 days were included as negative controls.

**Extended Data Fig. 5.**
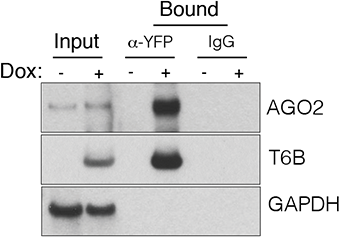
Total extracts from the colon of R26^T6B^ mice kept on doxycycline-containing diet for 1 week were immunoprecipitated using an anti-YFP antibody and probed with the indicated antibodies to measure the interaction between the T6B fusion protein and Ago2 *in vivo*. An anti-HA antibody was used to detect T6B.

**Extended Data Fig. 6.**
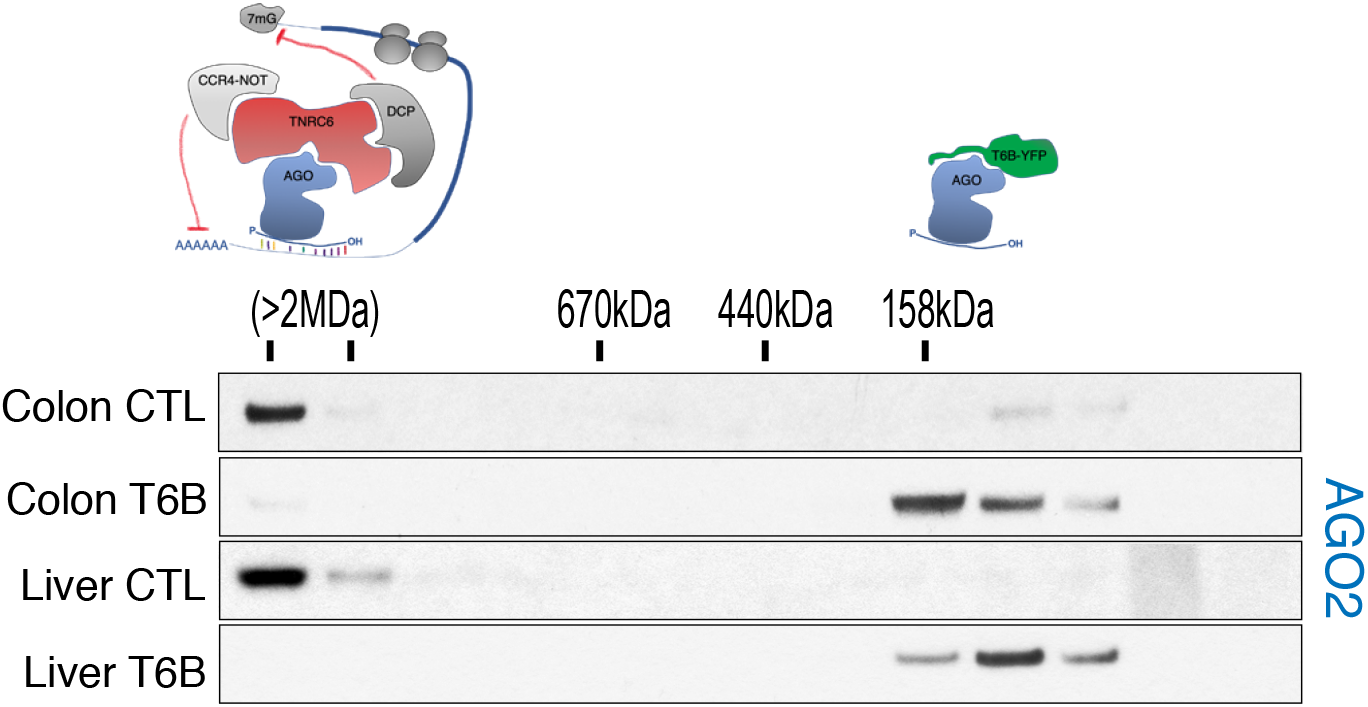
SEC fractionation followed by Western blotting of total extracts from the liver and large intestine of control and R26^T6B^ mice treated with doxycycline-containing chow for 7 days. The shift of AGO2 from high molecular weight to low molecular weight complexes confirms disruption of the miRISC.

**Extended Data Fig. 7.**
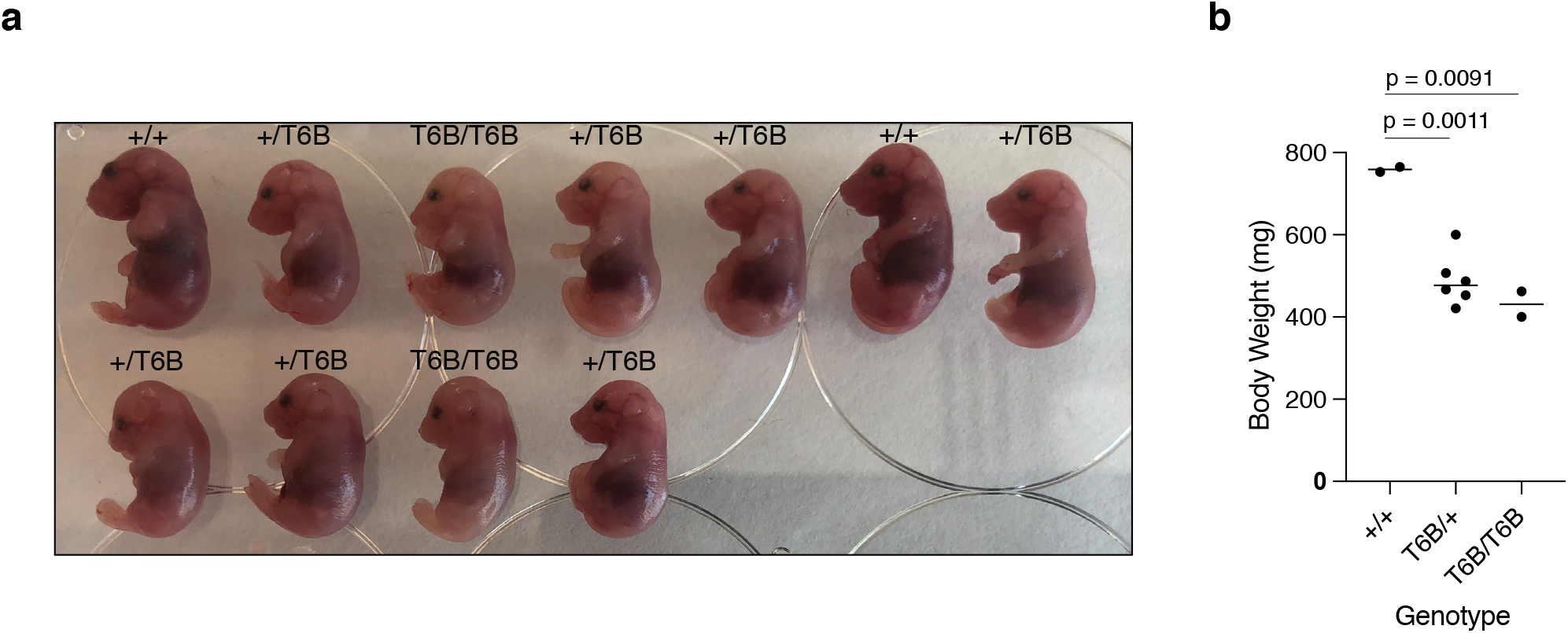
**(a)** Litter obtained by c-section from a pregnant R26^rtTA/rtTA^; ColA1^T6B/+^ female crossed to a R26^rtTA/rtTA^; ColA1^T6B/+^ male and maintained on doxycycline from d.p.c. 13.5 to d.p.c. 18.5. **(b)** Pups from (a) were weighted and genotyped and the results plotted. p-value: two-tailed unpaired t test.

**Extended Data Fig. 8.**
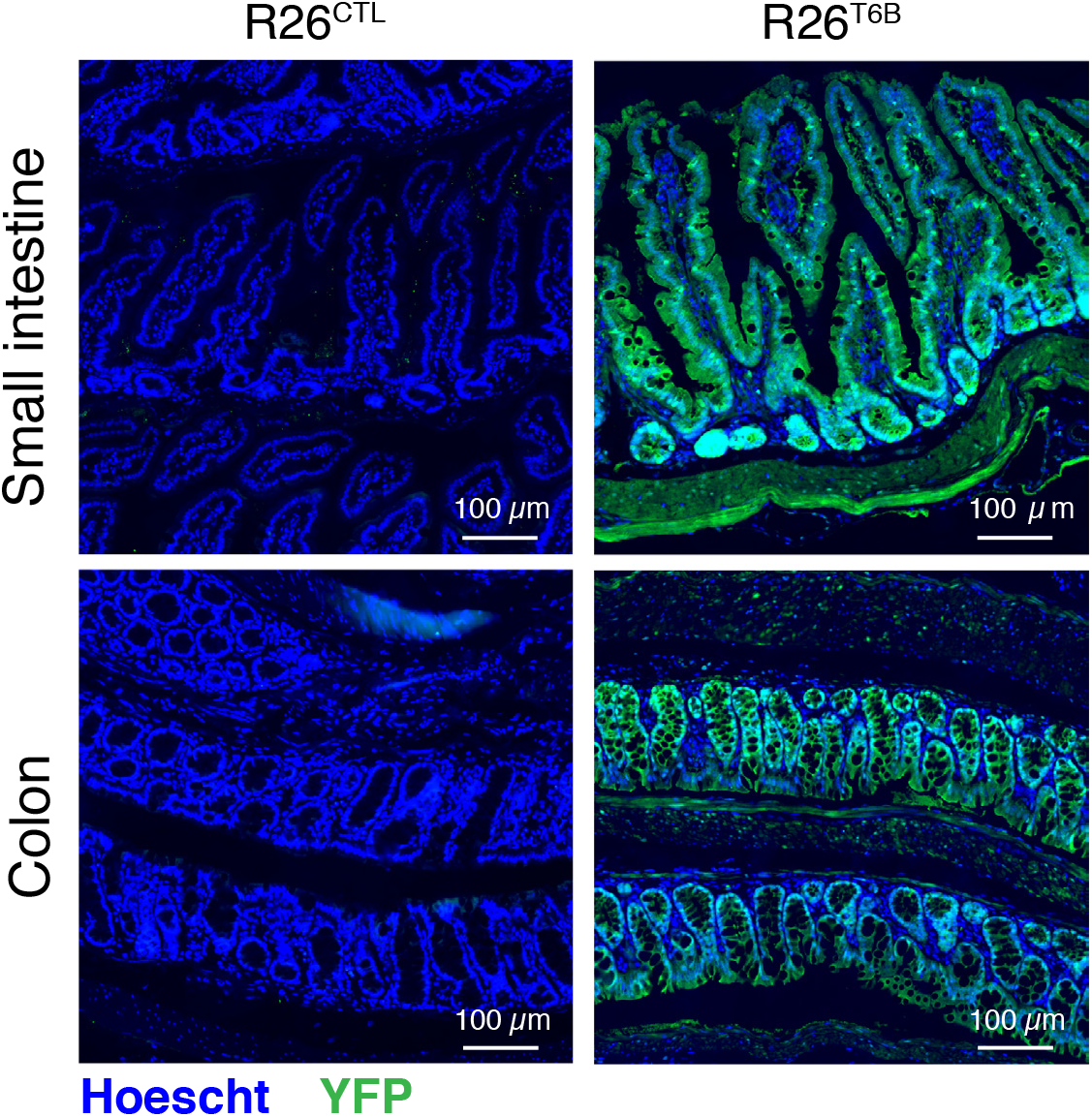
Immunofluorescence imaging of the small and large intestine of R26^T6B^ and R26^CTL^ mice kept on doxycycline diet for a month. An antibody against YFP was used to detect the T6B fusion protein.

**Extended Data Fig. 9.**
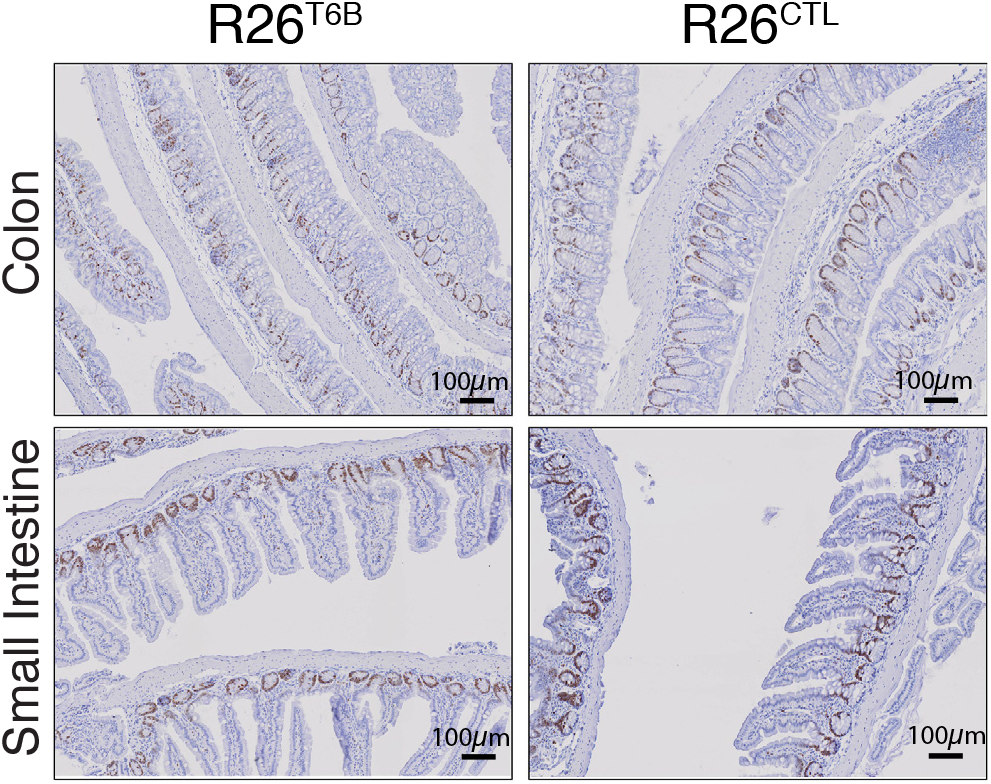
Sections from the colon and small intestine sections of R26^T6B^ and control mice kept on doxycycline-containing diet for 2 months were probed by IHC with an anti-Ki67 antibody.

**Extended Data Fig. 10.**
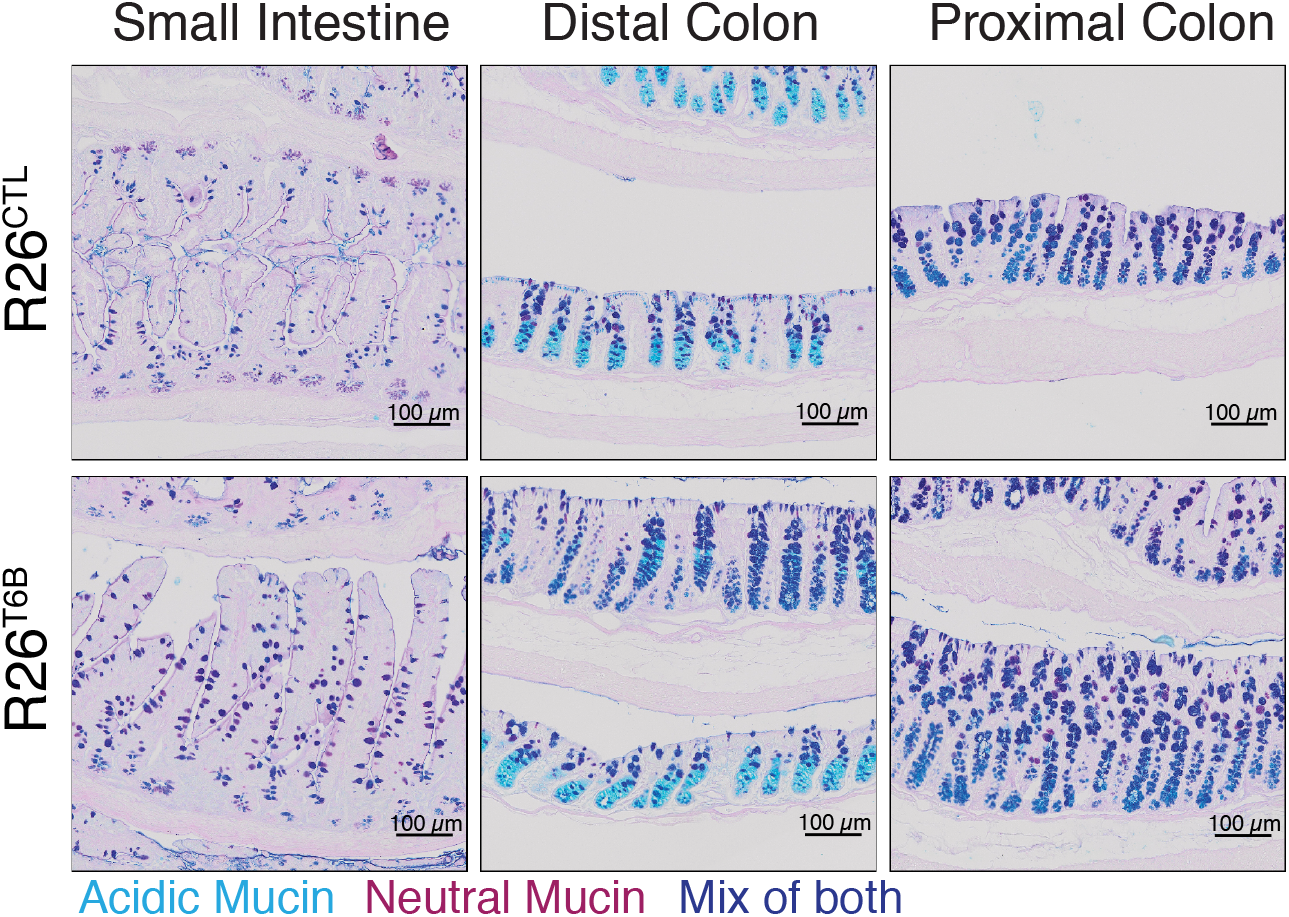
Detection of goblet cells by staining of acidic and neutral mucins in intestine sections from R26^T6B^ and control mice kept on doxycycline diet for 2 months. Neutral mucins are stained with periodic acid-Shiff whereas acidic mucins are stained with Alcian blue.

**Extended Data Fig. 11.**
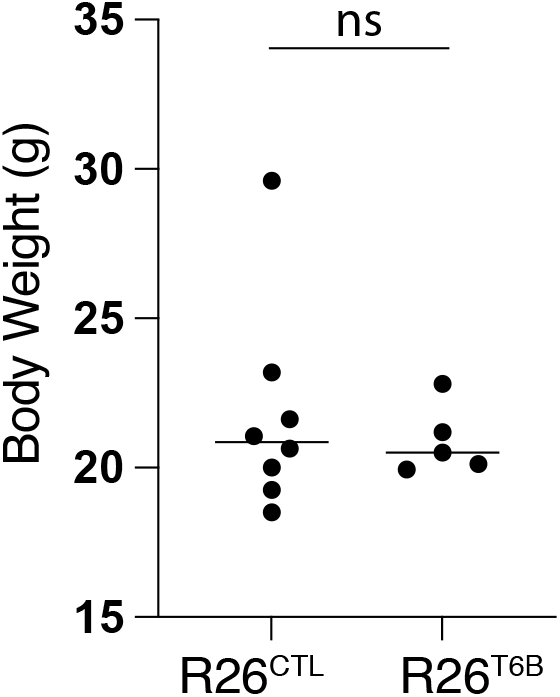
Body weight of R26^T6B^ (n = 5) and control (n = 8) female mice was assessed after 2 month-administration of doxycycline-containing chow. ns, not significant (p = 0. 6264, unpaired t test).

**Extended Data Fig. 12.**
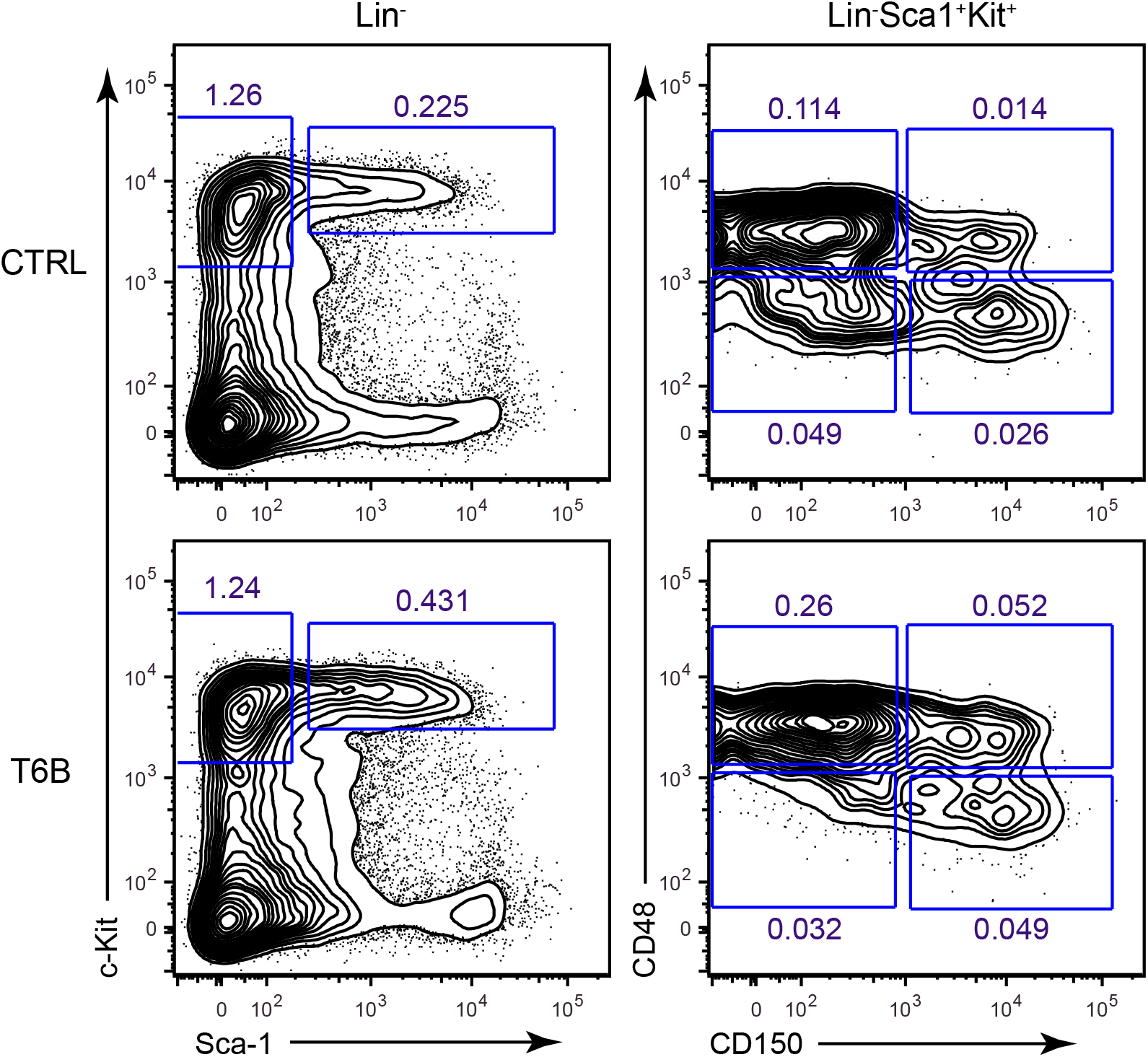
Representative flow cytometry plots showing the gating strategy for the identification of hematopoietic stem and progenitor cells from whole bone marrow harvested from R26^T6B^ and R26^CTL^ mice maintained on doxycycline diet for 3 weeks. LT-HSC: Lin- Kit+ Sca1+ CD150+ CD48-, ST-HSC: Lin- Kit+ Sca1+ CD150- CD48-, MPP2: Lin- Kit+ Sca1+ CD150+ CD48+, MPP3/4: Lin- Kit+ Sca1+ CD150- CD48+.

**Extended Data Fig. 13.**
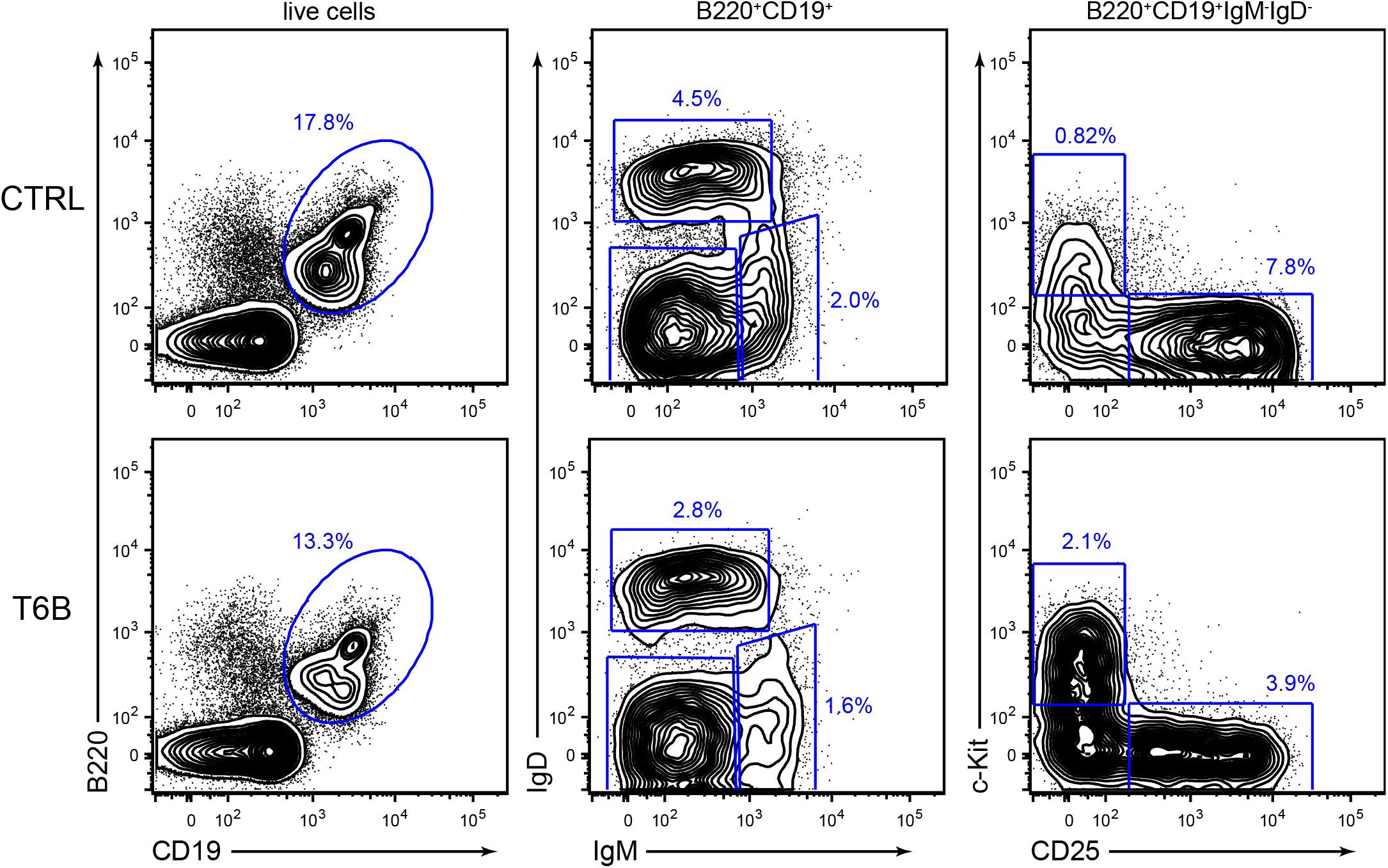
Representative flow cytometry plots showing the gating strategy for the identification of B cell lineage populations from whole bone marrow harvested from R26^T6B^ and R26^CTL^ mice maintained on doxycycline diet for 3 weeks. Pro-B: B220+CD19+IgD-IgM-CD25-Kit+, Pre-B: B220+CD19+IgD-IgM-CD25+, Imm B: B220+CD19+IgD-IgM+, Mat B: B220+CD19+IgD+IgM+/lo.

**Extended Data Fig. 14.**
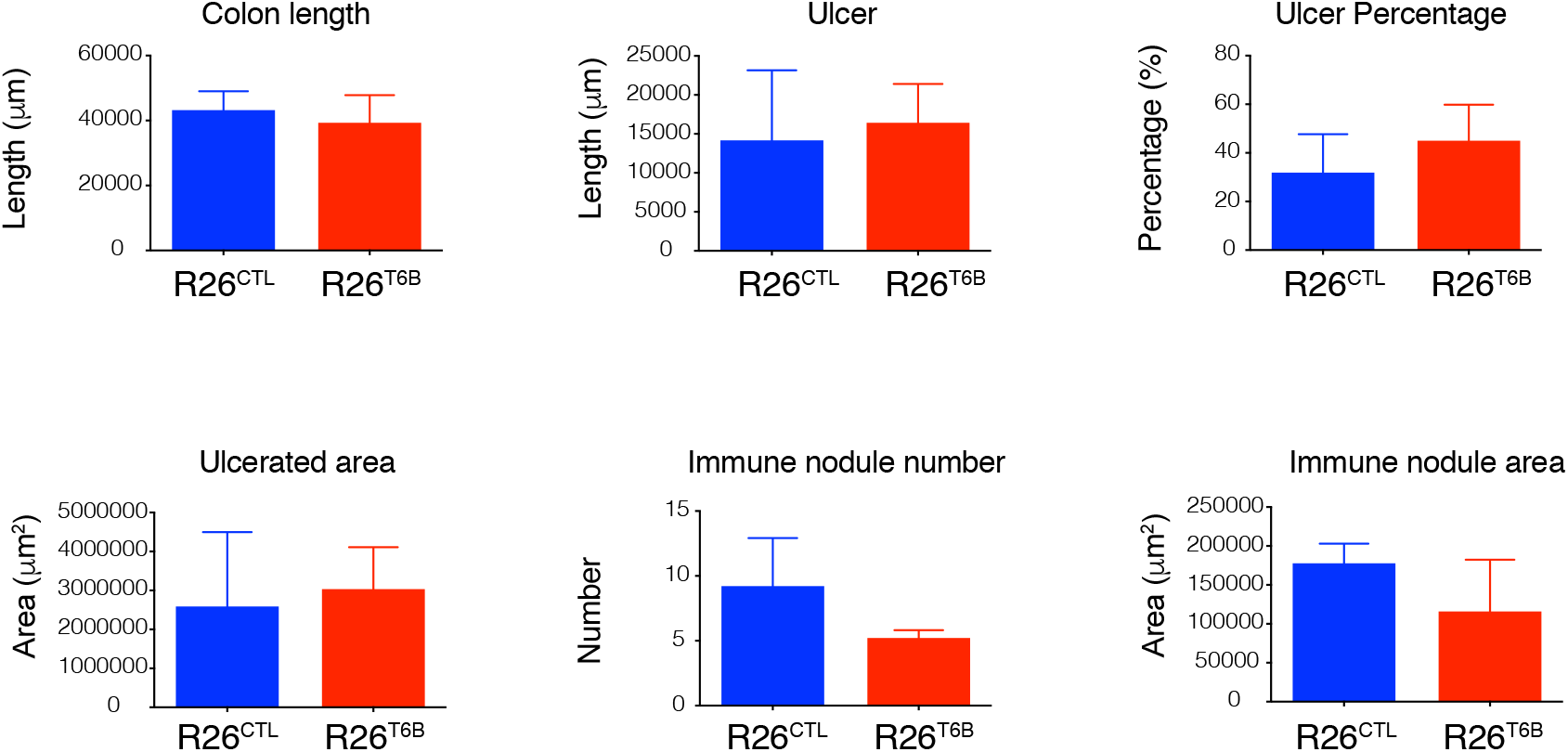
Bar plots showing measurement of colon length, aggregated length of ulcers, percentage of colon with ulcers, area of ulcers, number of immune nodules and the area of immune nodules performed on H&E longitudinal sections of colon from R26^CTL^ and R26^T6B^ mice 5 days post-DSS treatment. Measurements of these parameters were obtained using OMERO (https://www.openmicroscopy.org/omero/) and used to estimate the extent of damage and colitis induced by DSS treatment. Plots show that no significant differences between R26^CTL^ and R26^T6B^ mice were observed, suggesting that both groups experienced similar level of DSS-induced colitis.

**Extended Data Fig. 15.**
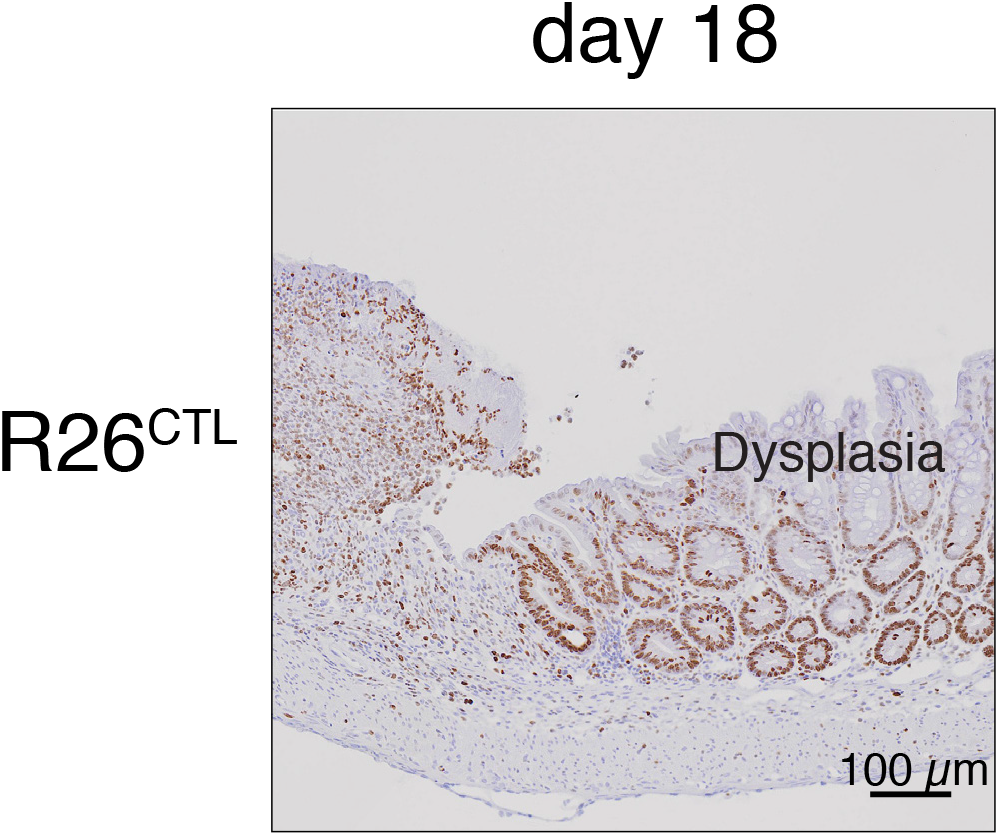
Representative immunohistochemistry image showing Ki67 signal in control mice (n = 3) 5 days after DSS treatment was discontinued. The presence of highly proliferating cells indicates residual dysplasia.

**Extended Data Fig. 16.**
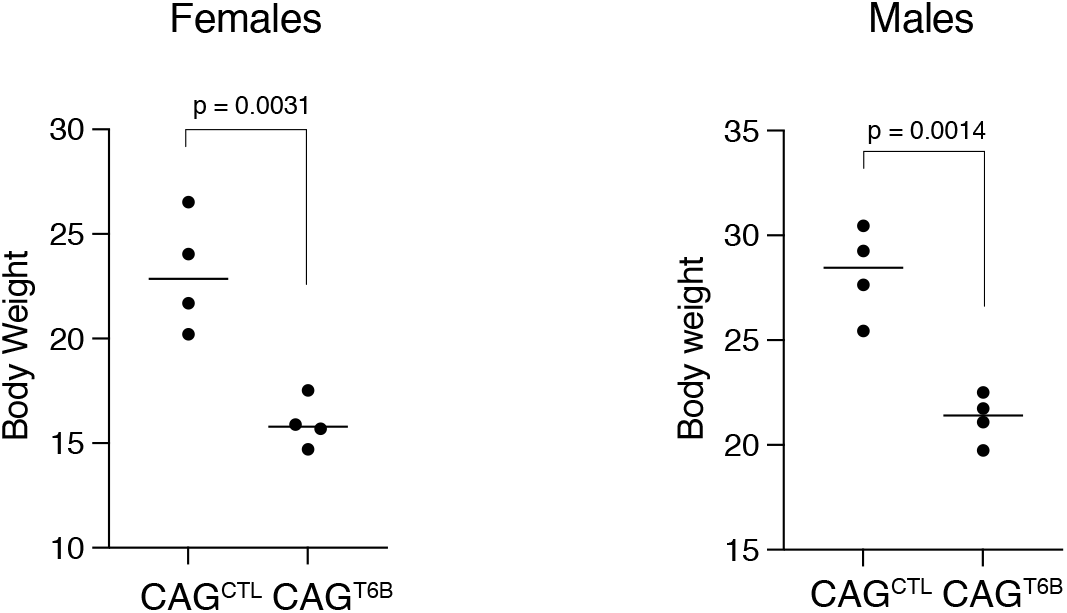
Body weight of CAG^T6B^ and control mice maintained on doxycycline for up to 45 days was assessed the day on which euthanasia was performed. n = 8 (4 females and 4 males) for each genotype (age and sex matched). Mice were kept on doxycycline diet throughout the duration of the experiment and control mice were euthanized at day 45. P-values: unpaired t-test.

**Extended Data Fig. 17.**
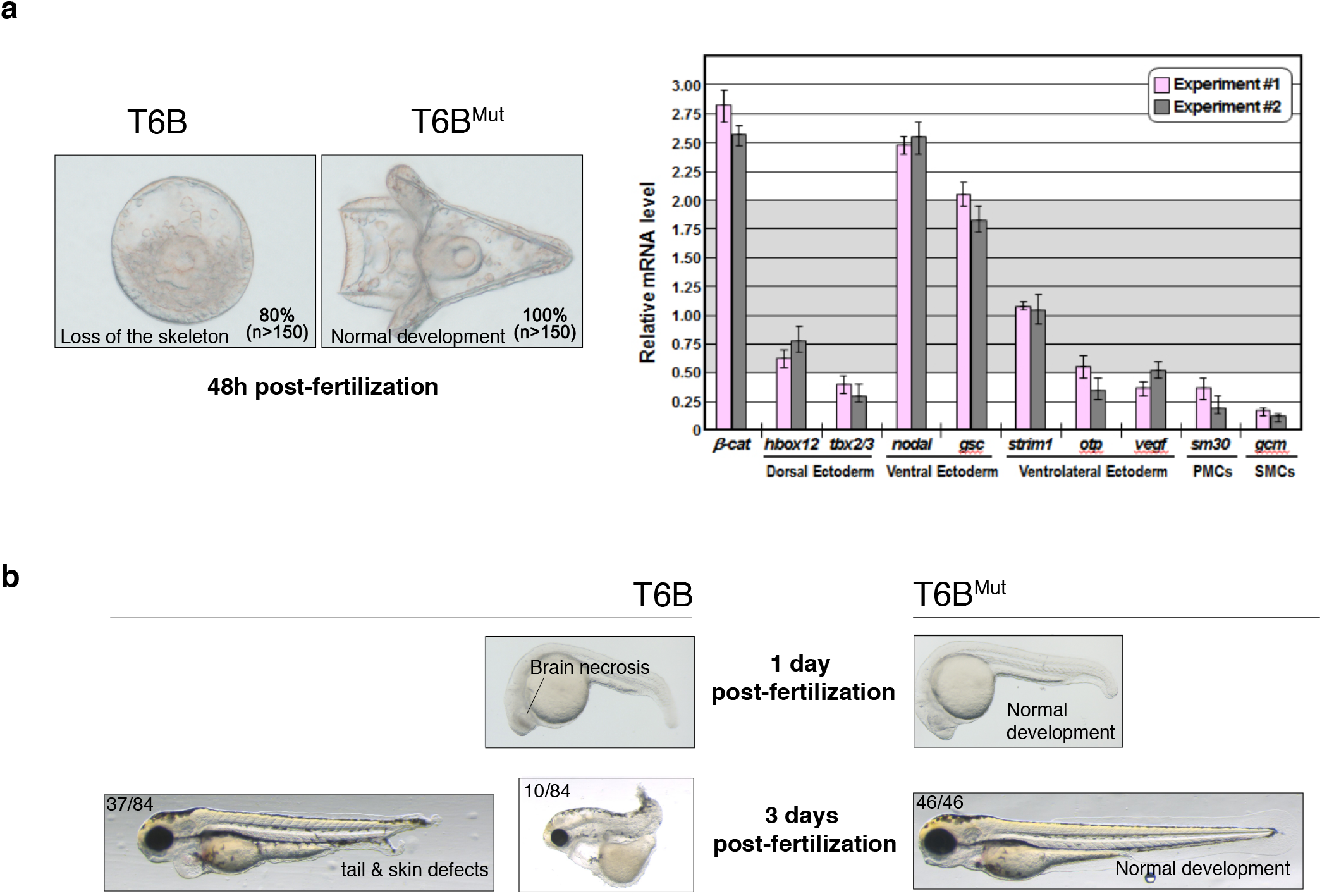
T6B blocks miRNA activity in sea urchins and zebrafish. **(a)** Left panel: Representative examples of Mediterranean sea urchin (*P. lividus*) zygotes injected with 1 pg of in vitro-transcribed mRNA coding for either T6B or T6B^Mut^ proteins and observed under DIC optics at 48 hours post-fertilization. Both embryos are oriented in a vegetal view. T6B-expressing embryos displayed severe developmental aberrations ranging from the failure to form a proper archenteron and skeletal structures, to overall delay in development and embryonic lethality. By contrast, control T6B^Mut^-expressing embryos observed at the same developmental stage went through embryogenesis normally and exhibited the characteristic easel-like shape of the echinoid pluteus larva. Right panel: quantitative PCR showing dysregulation of genes involved in the developmental gene regulatory network of the sea urchin^101^ upon T6B expression. **(b)** Zebrafish (*Danio rerio*) fertilized eggs were injected with 75 pg of in vitro-transcribed mRNA coding for either T6B or T6BMut fusion proteins. While T6B^Mut^-expressing embryos developed normally, the majority of T6B-expressing embryos under went severe developmental defects.

**Supplementary Table 1.** Complete blood counts (CBC) of whole blood from R26^T6B^ and R26^CTL^ mice taken after 3 weeks on doxycycline. Abbreviations areas follows: RBC = red blood cell count, HGB = hemoglobin, HCT = hematocrit, MCV = mean corpuscular volume, MCH = mean cell hemoglobin, RDW = red cell distribution width, RET = reticulocyte count, WBC = white blood cell count, PLT = platelet count.

|  |  | CTRL (n=5) | T6B (n=5) | *P* value |
| --- | --- | --- | --- | --- |
| RBC | 10<sup>6</sup>/μL | 10.8 ± 0.483 | 10.7 ± 0.292 | 0.9225 |
| HGB | g/dL | 16.4 ± 0.589 | 14.8 ± 0.277 | 0.0052 |
| HCT | % | 57.0 ± 1.74 | 52.0 ± 0.631 | 0.0037 |
| MCV | fL | 52.8 ± 1.33 | 48.6 ± 0.858 | 0.0041 |
| MCH | pg | 15.2 ± 0.2 | 13.9 ± 0.195 | 0.0002 |
| RDW | % | 22.8 ± 0.342 | 25.5 ± 0.716 | 0.0019 |
| RET | 10<sup>3</sup>/μL | 564 ± 68.5 | 569 ± 76.5 | 0.9273 |
| WBC | 10<sup>3</sup>/μL | 7.46 ± 1.16 | 9.16 ± 1.23 | 0.2608 |
| PLT | 10<sup>3</sup>/μL | 990 ± 290 | 1270 ± 154 | 0.2728 |

